# A germline PAF1 paralog complex ensures cell type-specific gene expression

**DOI:** 10.1101/2024.05.02.592153

**Authors:** Astrid Pold Vilstrup, Archica Gupta, Anna Jon Rasmussen, Anja Ebert, Sebastian Riedelbauch, Marie Vestergaard Lukassen, Rippei Hayashi, Peter Andersen

## Abstract

Animal germline development and fertility rely on paralogs of general transcription factors that recruit RNA polymerase II to ensure cell type-specific gene expression. It remains unclear whether gene expression processes downstream of such paralog-based transcription is distinct from that of canonical RNA polymerase II genes. In *Drosophila*, the testis-specific TBP-associated factors (tTAFs) activate over a thousand spermatocyte-specific gene promoters to enable meiosis and germ cell differentiation. Here, we show that efficient termination of tTAF-activated transcription relies on testis-specific paralogs of canonical Polymerase Associated Factor 1 Complex (PAF1C) proteins, which form a testis-specific PAF1C (tPAF). Consequently, tPAF mutants cause aberrant expression of hundreds of downstream genes due to read-in transcription. Furthermore, tPAF facilitates expression of Y-linked male fertility factor genes, and thus broadly maintains spermatocyte-specific gene expression. Consistently, tPAF is required for the segregation of meiotic chromosomes and male fertility. Supported by comparative *in vivo* protein interaction assays, we provide a mechanistic model for the functional divergence of tPAF and PAF1C and for transcription termination as a developmentally regulated process required for cell type-specific gene expression.

## INTRODUCTION

Across the tree of life, gene expression is facilitated by highly conserved core gene expression machineries, such as basal transcription factors, RNA polymerase complexes, RNA export factors, and ribosomes. These core gene expression machineries are generally not considered regulatory and rather facilitate gene expression globally and across all cell types in multicellular organisms. Cell type identity in multicellular organisms is instead controlled by a separate layer of gene regulatory proteins, many of which recognize target genes or RNA in a sequence-specific manner, such as is the case for sequence-specific transcription factors, RNA-binding proteins, and microRNAs^1–3^. While this broad textbook generalization overall aligns with our current knowledge of gene expression, examples of tissue- and cell type-specific variants of core gene expression machineries have been identified in most animal model organisms, including mice, frogs, zebrafish, and fruit flies (*Drosophila melanogaster*). These machineries include paralogs of basal transcription factors^4–12^, RNA exporters^13,14^, ribosomal proteins^15–17^, and proteasome subunits^18^. The molecular function and developmental impact of such core gene expression machinery variants, however, remain underexplored.

Notably, most identified paralogs of core gene expression machineries are expressed specifically in the germline tissue of the male (testis) and female (ovary) animals. While some of the identified paralogs are likely non-essential pseudogenes, several have been found to be required for proper germline development and animal fertility^4,7,8,11–14,19–22^. The lack of tractable *in vitro* systems for animal germline biology has limited detailed molecular characterization with only a few paralogs having been characterized at the molecular level. Such examples include the mouse TATA box Binding Protein (Tbp) paralog TBPL2/TRF3, which replaces the general transcription factor TBP in growing oocytes and mediates transcription of oocyte-expressed genes^6^ and the *Drosophila* TFIIA paralog, Moonshiner, which enables transcription of transposon-silencing piRNA precursors from otherwise silent heterochromatin^4^. Germline-specific paralogs of core gene expression machinery thereby facilitate germline development and animal fertility by enabling alternative modes of gene expression.

Germ cell differentiation from proliferating spermatogonia to meiotic spermatocytes comprises one of the most substantial genome reprogramming events known in animals and entails thousands of testis-specific genes being activated, shut down, or expressed in alternative isoforms^23,24^. In *Drosophila melanogaster*, this dramatic genome regulation relies on testis-specific paralogs of core gene regulatory factors, including bromodomain proteins (tBRDs)^25,26^, the dREAM complex (tMAC)^27^, and the TFIID complex (tTAF)^28,29^. Consistently, mutation of the tTAF genes, *cannonball* (*can* / *TAF5L*), *no hitter* (*nht* / *TAF4L*), *meiotic arrest* (*mia* / *TAF6L*) and *spermatocyte arrest* (*sa* / *TAF8L*) all lead to a block of meiotic cell cycle progression and complete infertility in male flies^28,30^. Mechanistically, tTAF and tMAC proteins are thought to regulate gene expression through direct association with the promoter regions of the target genes^23^. However, it remains unclear how these complexes are functionally connected and whether they couple to downstream non-canonical gene expression regulators to enable germline development.

Here we identify a variant Polymerase Associated Factor 1 Complex (PAF1C) as an essential transcriptional regulator required for *Drosophila* male germline development. The testis-specific PAF1C (named tPAF) is composed of paralog proteins of each of the four canonical PAF1C core proteins as well as the PAF1C-associated protein, Ski8. tPAF is expressed specifically during the spermatocyte stage of male germline development during which meiotic defects in chromosome separation arise in tPAF mutant flies. Mechanistically, tPAF physically associates with the tBRD and tTAF proteins and functions to ensure efficient transcription termination specifically at genes that require tTAF for their activation. In addition, the tPAF subunit Cdc73L is largely dispensable for tPAF-mediated transcriptional regulation but is required for the nucleolar accumulation of both tPAF and tTAF. tPAF thus physically and functionally connects germline-specific variant core genome regulation complexes and reveals how the fundamental reprogramming during animal germline development is enabled by the concerted function of paralog-based variants of core gene expression regulators.

## RESULTS

### Paralogs of the PAF1 complex are required for male fertility in *D. melanogaster*

To probe the existence of functional connections between paralog-based genome regulators in the fly male germline model system, we searched for novel tTAF protein associations using *in vivo* proximity labelling coupled with mass-spectrometry analyses. From confocal microscopy of testis from flies harboring the lowly expressed TurboID ligase–anti-GFP nanobody fusion protein^31^ and a GFP-tagged *spermatocyte arrest (sa)* tTAF allele, we observed a marked accumulation of biotinylation in spermatocyte nucleoli, overlapping GFP-tTAF protein (**Figure S1A**), thereby indicating high specificity and efficiency of the proximity labelling. Mass-spectrometry analyses of biotinylated proteins from lysates of Sa-GFP revealed strong enrichment relative to control testis of basal transcription factor proteins (**Figure 1A**, light blue data points), including known tTAF subunits, as well as CG14930, a previously unrecognized paralog of the TFIID subunit Taf9 (**Figure S1B**) that we therefore name *Taf9L*. We note that the known tTAF protein Taf12L becomes undetectable by imaging when used as proximity labelling bait (**Figure S1C**) and it is therefore likely missing in the Sa proximity labeling data due to biotin-related degradation. BLAST searches of the other highly enriched uncharacterized proteins revealed two proteins with strong homology to the Paf1/atms and Leo1/Atu subunits of the Polymerase associated factor 1 complex (PAF1C), an essential and highly conserved eukaryotic RNA polymerase II regulatory complex^32^. We therefore named the two genes based on this homology: *CG12674* as *Paf1/atms-like* (*Paf1L*) and *CG10887* as *Leo1/Atu-like* (*Leo1L*). Reverse BLAST searches for the two other core PAF1C subunits, *Cdc73/hyx* and *Ctr9*, identified *CG6220* (renamed *Cdc73L*) and *CG9899* (renamed *Ctr9L*) to reveal a complete set of PAF1C core subunit paralogs enriched by Sa-GFP proximity-labeling (**Figure 1A**, orange data points).

**FIGURE 1.**
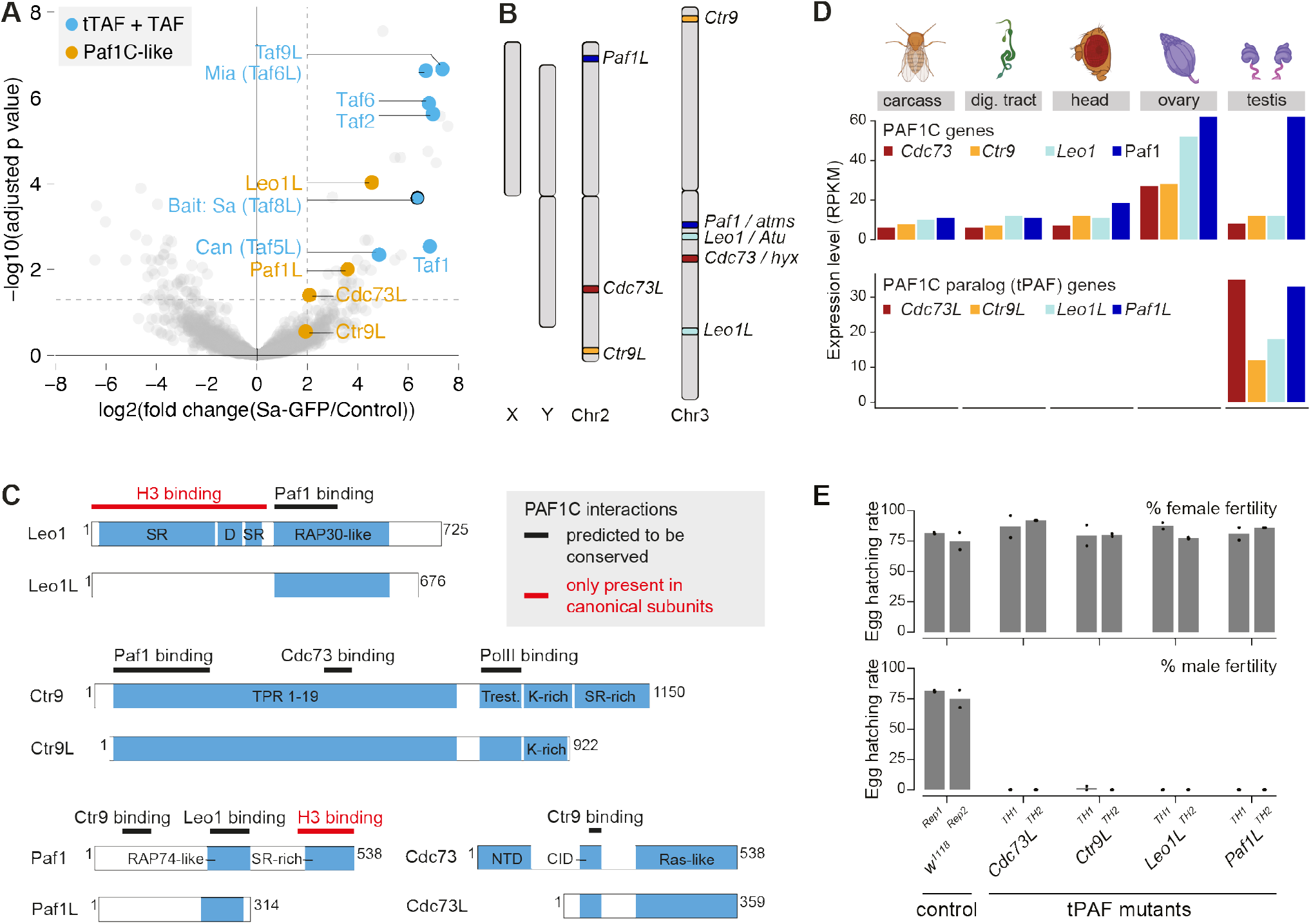
Paralogs of the Paf1 complex are required for male fertility in D. melanogaster. (A) Enrichment values and corresponding statistical significance levels for proteins associated with the *Spermatocyte arrest* (*Sa*; tTAF) bait protein as detected by proximity labeling mass-spectrometry. Blue highlights indicate basal transcription factors, including tTAF. Orange highlights indicate identified PAF1C-like proteins. (B) Schematic of the chromosomal position of PAF1C genes and the identified PAF1C gene paralogs. (C) Schematic of the aligned PAF1C and tPAF genes with the main protein domains as well as known interaction surfaces from PAF1C indicated. (D) mRNA levels of PAF1C and PAF1C paralog (tPAF) genes in selected adult tissues (RNAseq data from modENCODE, ref. ^90^), RPKM = Reads Per Kilobase per Million mapped reads. (E) Bar plot showing the mean embryo hatching rate in percent of eggs laid by control (*w*^*1118*^) and tPAF mutant flies. Individual data points from biological replicates (n = 2) are shown as black dots.

The PAF1C paralog genes originated 50-100 million years ago (**Figure S2A-B**), likely through retroduplication of the canonical PAF1C genes as indicated by their comparatively low number of introns and spread across the genome (**Figure 1B**). Since then, the PAF1C paralog genes have undergone notable sequence divergence, including loss of several protein domains important for canonical PAF1C function (**Figure 1C**). The paralog genes have, however, retained the homologous regions known to facilitate PAF1C complex formation (**Figure 1C**), indicating maintenance of some functionality. While canonical PAF1C is expressed in all tissues, as expected from its role in eukaryotic RNA polymerase II regulation, the PAF1C paralog genes are expressed exclusively in testis (**Figure 1D**) and we refer to these as tPAF for testis-specific PAF1C. To assay the putative function of the tPAF genes we generated flies harboring CRISPR/Cas9-induced frame-shift mutations (**Figure S2C**). Transheterozygous mutants of all four tPAF genes are viable but display full sterility in males, consistent with an essential function in testis genome regulation (**Figure 1E**). A complementary short hairpin RNA (shRNA)-based depletion approach, using a germline-specific knock-down driver, validated the sterility phenotype and further supported that tPAF functions in the testis germline cells (**Figure S3A**). In sum, we conclude that tPAF genes, despite pronounced sequence divergence, have likely maintained genome regulatory function and evolved to become essential for male fertility.

### tPAF mutant males exhibit meiotic defects

To elucidate the role of tPAF genes in male fertility, we investigated potential morphological anomalies occurring during spermatogenesis in tPAF gene mutants. During spermiogenesis, the final stage of spermatogenesis, spermatids transform into spermatozoa. This transformation involves a series of events, including the initial elongation of nebenkern (mitochondrial derivatives) along the axoneme, subsequent nuclear elongation, histone-to-protamine exchange, and finally, spermatid individualization **(Figure 2A)**. Previous studies have demonstrated that mutants of the tMAC and tTAF complexes exhibit a complete meiotic arrest at the pre-meiotic spermatocyte stage, leading to the absence of post-meiotic germline cells^28,30,33^. We were able to recapitulate this meiotic arrest phenotype using a mutant of the tMAC gene *always early (aly)*, which was evident from the absence of post-meiotic germline cells. However, tPAF mutant alleles did not display such a meiotic arrest as post-meiotic germline cells were still observed **(Figure 2B, Figure S4A)**. To understand where in germline development tPAF mutant sterility arises, we first investigated spermatid individualization. In contrast to the distinctive morphology in control samples with aligned elongated bundles of needle-like spermatid nuclei trailed by actin cones **(Figure 2C, left)**, tPAF mutants exhibited pronounced deficiencies in spermatid maturation characterized by rounded nuclei, ‘lagging’ actin cones, and a lack of polarization as marked by Orb protein **(Figure 2C-D, Figure S4B**,**D-E)**. Furthermore, we observed the association of transition proteins and protamines with the disorganized spermatid nuclei in Paf1L mutants, indicating that DNA compaction itself remained unaffected **(Figure S5)**. To understand whether the defects resulting in faulty spermatid individualization arise before or after completion of meiosis in spermatocytes, we next characterized early spermatids at the ‘nebenkern/onion’ stage, where aberrant nuclear morphology is a known hallmark of meiotic defects in the preceding spermatocyte stage^34^. In control testes, the majority of spermatids exhibited nuclei and nebenkern structures that were spherical, of equal size, and occurred in pairs (**Figure 2E, left)**. By contrast, tPAF mutant testes displayed pronounced abnormalities in the majority of spermatid cysts **(Figure 2E-F; Figure S4C**,**F)**. Abnormalities include irregular shapes, enlarged nebenkern, and multinucleation – defects which strongly suggest failure in cytokinesis during meiosis^34^.

**FIGURE 2.**
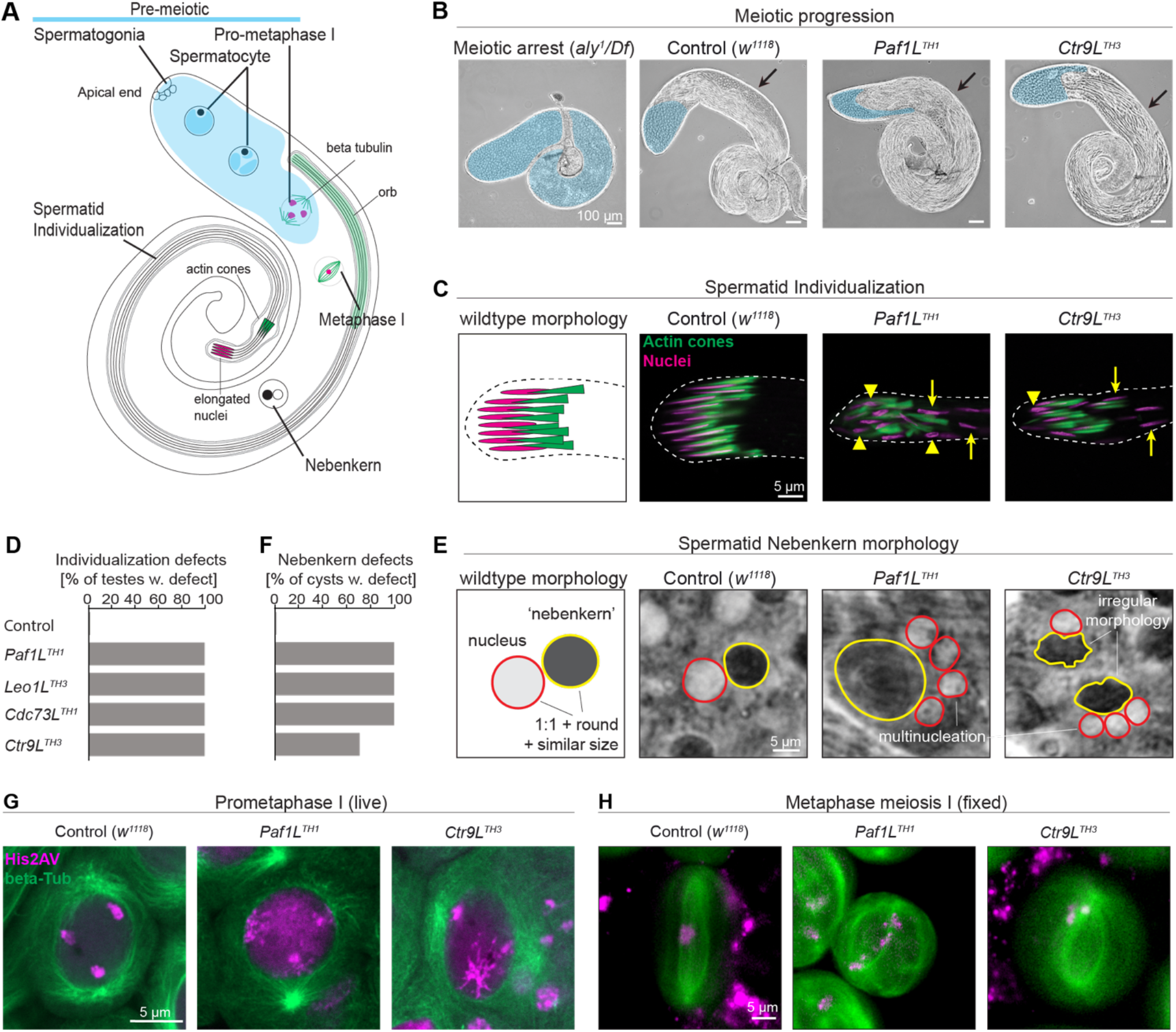
tPAF mutant male gametogenesis shows defects originating from abnormal meiosis. (A) Illustration of spermatogenesis in male Drosophila. Shaded cyan region: pre-meiotic stages, including spermatogonia and spermatocytes. (B) Phase contrast images comparing wild type to meiotic arrest control (*aly*^*1/Df*^) and tPAF mutants. Cyan region: pre-meiotic stages. Arrow: post-meiotic regions. (C) Immunostaining images of wild-type controls and tPAF mutants showing abnormalities during individualization. Magenta: spermatid nuclei visualized by H2Av-RFP. Green: spermatid F-actin cones, stained by FITC-phalloidin. Spermatid individualization defects are highlighted: disruption of the characteristic needle-like structure of the nuclei (arrowheads), and a lack of coordinated localisation of the nuclei (arrows). (D and F) Bar charts showing the number of defective elongated spermatids (D) and defective nebenkern stage spermatids (F) in WT and tPAF mutants (n=10, representative of independent experiments). Defective elongated spermatids are defined as spermatids that carry 5 or more nuclei with described defects. (E) Phase contrast images of the nebenkern stage spermatids from the control and tPAF mutant testes, showing an array of defects, including enlarged or irregularly shaped nebenkern surrounded by multiple small nuclei. (G and H) Fluorescent microscopy imaging of spermatocytes at pro-metaphase I (G, live imaging) and metaphase I (H, fixed cells), comparing wild type to tPAF mutants. Magenta: chromosomes visualized by H2Av-RFP. Green: microtubules visualized by beta-Tubulin-GFP. Pro-metaphase I and metaphase I identification are based on the formation of astral fibers around the nucleus and the spindle formation, respectively. Images are representative independent experiments (n = 10 (G); n = 5 (H)). Images in the same panel are on the same scale (see scale bars).

To understand the possible meiotic defects in tPAF mutants we next investigated meiosis using live imaging. In control spermatocytes, pro-metaphase is characterized by condensed chromosomes occupying three distinct territories at the periphery of the cell— two autosomal territories and an X-Y territory^35^ (**Figure 2G, left panel**). tPAF mutants, however, display a lack of well-defined territories, with His2Av-RFP diffusely present in the center of the cell (**Figure 2G, S6A**). This suggests that tPAF mutants exhibit incomplete chromosomal condensation by pro-metaphase. During metaphase I, these condensed chromosomes migrate to the central part of the cell, forming a distinct mass. Concurrently, the nuclear membrane breaks down, and astral fibers penetrate into the nucleus, securing the condensed chromosomes^35^ (**Figure 2H, left panel**). Though astral fiber formation is observed, tPAF mutants lack a central mass during metaphase I (**Figure 2H, S6B**), likely leading to unequal chromosomal division during anaphase I. In sum, tPAF mutants exhibit aberrant chromosome movement during meiosis, which likely originates from faulty chromosome condensation in pre-meiotic spermatocytes. Together, these observations suggest that tPAF functions to enable proper spermatocyte maturation and differentiation.

### tPAF forms a complex *in vivo* in spermatocytes that is distinct from canonical PAF1C

To characterize the molecular function of the tPAF genes, we next generated endogenously tagged alleles of all four tPAF genes as well as of three canonical PAF1C genes by integrating a localization and purification (LAP)-tag consisting of triple FLAG and V5 tags as well as GFP. LAP-tag integration using CRISPR/Cas9 resulted in fully viable and fertile flies showing the expected fusion protein expression in testis lysates (**Figure 3A; Figure S8A**). All tagged tPAF proteins showed a similar localization with strong expression specifically in spermatocytes (**Figure 3B; Figure S7A-B**), consistent with the proposed functional role during spermatocyte differentiation and meiosis. On the other hand, canonical PAF1C proteins are expressed already in mitotic spermatogonia, as well as in spermatocytes and the surrounding somatic muscle sheath cells (**Figure 3C; Figure S8B**).

**FIGURE 3.**
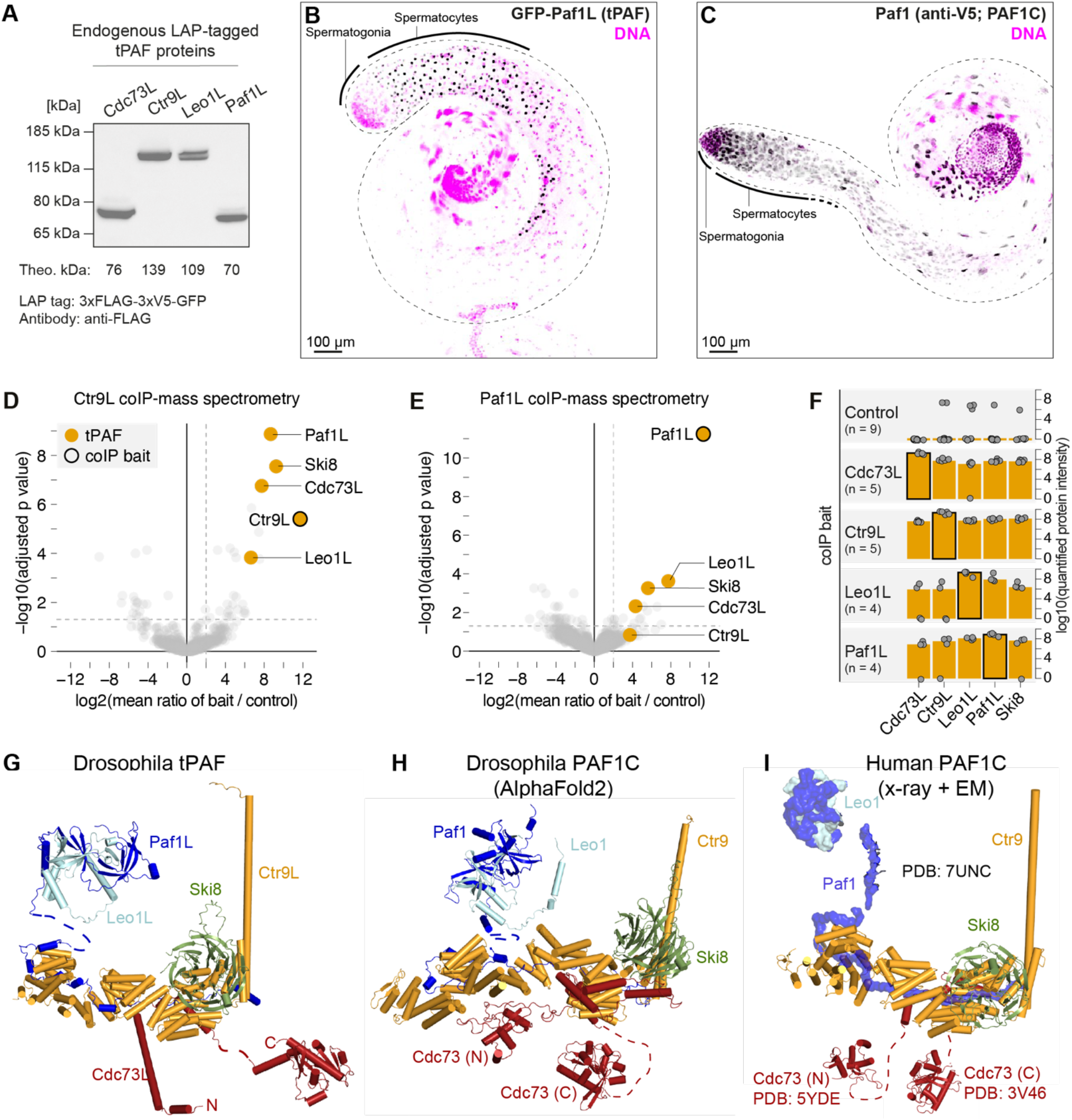
tPAF forms a complex in vivo in spermatocytes that is distinct from canonical PAF1C. (A) Western blot detection of endogenously tagged tPAF proteins using anti-FLAG antibodies. (B-C) Confocal images showing the GFP signal in testis tissue from endogenously tagged Paf1L tPAF (B) or Paf1 PAF1C (C) proteins with DAPI-stained DNA shown in magenta. (D-E) Enrichment values and corresponding statistical significance levels for proteins co-immunopurified (coIP) with endogenously tagged Ctr9L (D) or Paf1L (E) proteins as detected by mass-spectrometry (biological replicates, n, are indicated in F). (F) Bar plot showing the median quantified tPAF protein intensity of control and tPAF protein coIP-mass-spectrometry biological replicates. The individual replicate protein intensities are plotted with grey circles. (G-H) Predicted structure model of tPAF (G) and Drosophila PAF1C (H) complexes based on Alphafold2. Dashed lines indicate manually positioned structured domains. (I) Structural model of human PAF1C based on x-ray crystallography and cryo-electron microscopy related to the indicated PDB entries.

We next used the 3xFLAG epitope of the tPAF fusion proteins to investigate whether tPAF proteins are integrated into complexes *in vivo* by co-immunoprecipitation (co-IP) coupled to protein identification by mass spectrometry. We produced protein lysates by cryo-grinding whole male flies, thereby creating a highly complex protein lysate for stringent co-IP assays. co-IP of Ctr9L and Paf1L showed marked enrichment of the other three tPAF proteins (**Figure 3D-F)**. Given the low number of spermatocytes in the whole-male lysate, these results strongly indicate that (i) tPAF proteins form a complex *in vivo* and (ii) the tPAF complex forms exclusively from tPAF proteins without cross-interaction with the core canonical PAF1C proteins. Leo1L co-IP showed a notably stronger association with Paf1L than with the other tPAF proteins (**Figure S9A)** and Leo1L protein levels were comparably low in the Cdc73L co-IP (**Figure S9B)**, indicating that Leo1L may form a sub-complex with Paf1L, analogous to what has been observed for the canonical PAF1C^36^. In addition to the core tPAF proteins, Ski8/CG3909, was strongly enriched in all tPAF protein co-IP experiments (**Figure 3D-F; Figure S9A-B**). Ski8 is a member of the conserved cytoplasmic RNA decay cofactor, the *Super Killer* (SKI) complex conserved from yeast to flies and humans^37,38^. In mammals, but not in yeast, Ski8 is also part of canonical PAF1C^39^. Conversely, yeast PAF1C contains the protein Rtf1^40,41^, which is not part of human PAF1C. While we identified a testis-specifically expressed Rtf1 paralog with a similar evolutionary origin as the tPAF genes (**Figure S10A-C**), Rtf1L mutants showed no fertility defects and endogenously LAP-tagged Rtf1L protein neither co-localized with nor co-IP’ed tPAF proteins (**Figure S10D-H**).

Structure modeling of tPAF using Alphafold2^42^ predicts that it forms a complex with a very similar overall structure to that modeled for canonical Drosophila PAF1C as well as to the experimentally determined human PAF1C structure (**Figure 3G-I, S11A-D)**. We therefore conclude that tPAF forms a protein complex consisting of the paralog proteins Paf1L, Leo1L, Cdc73L, and Ctr9L as well as Ski8 *in vivo*, constituting a protein composition similar to mammalian canonical PAF1C.

### tPAF is required to maintain spermatocyte-specific gene expression

The prominent sub-nuclear localization of tPAF is similar to that described for tTAF and tBRD proteins, which are known to accumulate in spermatocyte nucleoli^25,29,43^. Indeed, co-visualization of LAP-tagged tPAF proteins with the nucleolar protein Fibrillarin revealed a clear nucleolar accumulation of all tPAF proteins in the granular component nucleolar subregion outside the Fibrillarin-dense areas (**Figure 4A; Figure S12A**), overlapping with the localization of the GFP-tagged Sa (tTAF) protein (**Figure S12B**). tPAF initially accumulates in the nucleolus as spermatogonia differentiate into spermatocytes and then remains in the nucleolar region even after Fibrillarin has left the nucleoli. The nucleoli disappear before cells enter meiosis where the tPAF proteins are detectable only as small DNA-associated foci (**Figure S13A-B**). Of note, a diffuse but notable tPAF protein signal was also observed over the nuclear chromosome territories **(Figure 4A, right panels)**, suggesting that tPAF, similar to tTAF proteins^29^, is also present on the main nuclear chromosomes and may function in the nucleoplasm too.

**FIGURE 4.**
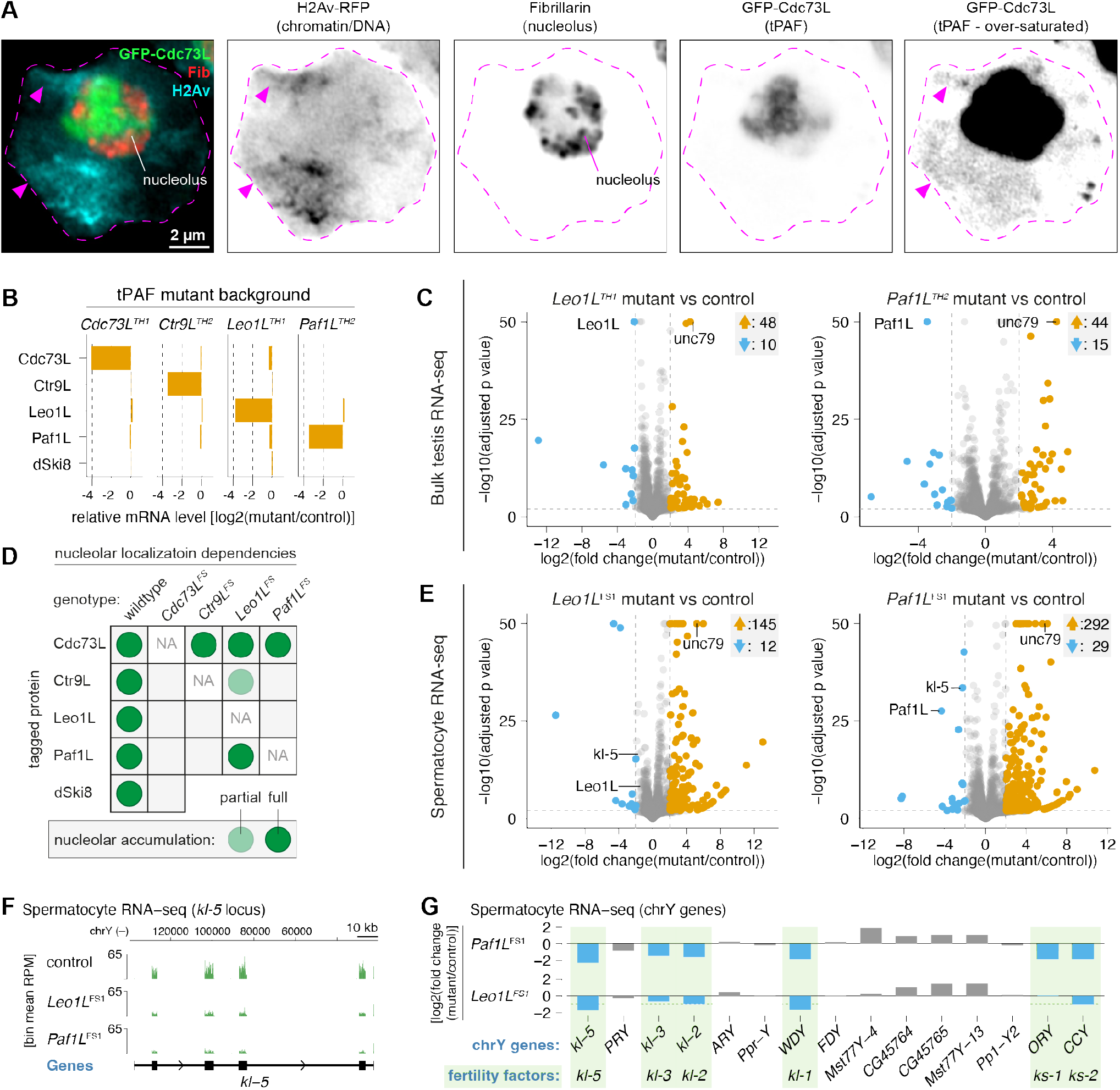
Spermatocyte-specific gene expression depends on tPAF. (A) Confocal microscopy images showing the localization of GFP-tagged Cdc73L, Fibrillarin (Fib, anti-Fib immunoflourence staining), and chromatin visualized by H2Av-RFP. Dashed magenta line: nuclear border. Magenta arrowheads: position of major chromosome territories. (B) Bar plots showing the levels of tPAF gene mRNA based on bulk testis RNAseq from various tPAF mutants relative to control. (C) Enrichment values and corresponding statistical significance levels for mRNAs quantification based on bulk testis RNAseq in biological replicates of *Leo1L* (left panel) or *Paf1L* (right panel) tPAF mutants (n = 3) compared to control (n = 6) flies. Orange and blue arrows: the number of genes categorized as upregulated and downregulated, respectively. (D) Summary schematic of the identified interdependencies for nucleolar accumulation of tPAF proteins based on the data shown in Figure S16. (E) Enrichment values and corresponding statistical significance levels for mRNAs as quantified by RNAseq from FACS-sorted Cdc73L-GFP-positive spermatocytes in biological replicates of *Leo1L* (left panel) or *Paf1L* (right panel) tPAF mutant or control flies (n = 3). (F) Genome tracks showing RNA-seq coverage around the *kl-5* gene for the indicated genotypes. Bin mean RPM: mean reads per million mapped reads (RPM) per genome region bin. (G) Bar plots showing the log2 fold change in mRNA levels of Y-linked genes, comparing tPAF mutant and control genotypes. Blue bars: mRNA level at least 50 % decreased in any tPAF mutant sample compared to control (Figure S14E). Green highlights: male fertility factor genes.

Since canonical PAF1C is known to function in transcriptional regulation, we next investigated whether the tPAF complex might also serve such functions. To this end, we performed transcriptome profiling of total testis RNA derived from control and mutants of each tPAF subunit. All tPAF mutant samples showed a pronounced depletion of mRNA from the mutated gene **(Figure 4B)**. Consistent with a function in gene regulation, differential gene expression analysis showed that mutants of *Paf1L* and *Leo1L* result in notable transcriptome changes **(Figure 4C)** although at modest levels compared to the dramatic changes in mutants of the *cannonball (can)* (tTAF) and *cookie monster (comr)* (tMAC) subunit (**Figure S14A-B**). By contrast, gene expression in *Cdc73L* and *Ctr9L* mutants was not strongly affected **(Figure S14C-D)**, suggesting that tPAF-mediated gene regulation relies mainly on Paf1L and Leo1L.

To better understand such subfunctionalizations within the tPAF complex, we assayed the localization of all endogenously tagged tPAF subunits in the mutant background of each of the other tPAF genes. Consistent with Cdc73L serving other functions, potentially related to the nucleoli, we found that Cdc73L independently localizes to the nucleolus and is required for nucleolar accumulation of all other tPAF proteins (**Figure 4D; Figure S15A-B**). In addition, Paf1L and Ctr9L showed interdependence, while Leo1L was largely dispensable for nucleolar localization of the remaining tPAF proteins. Although no notable GFP signal was visible in localization-disrupting mutants, western blotting analysis demonstrated that tPAF proteins remained expressed, indicating dispersal upon loss of nucleolar localization (**Figure S15C**). We therefore conclude that tPAF harbors sub-functionalization with Cdc73L being largely dispensable for tPAF-mediated gene regulation but required for tPAF accumulation in nucleoli.

tPAF proteins are specifically expressed in spermatocytes and bulk tissue transcriptome analysis may therefore mask the true regulatory phenotype. We therefore took advantage of the observed sub-functionalization and used the detection of Cdc73L-GFP to purify tPAF-expressing spermatocyte cells in control as well as *Leo1L* and *Paf1L* mutants and performed transcriptome profiling (**Figure S16A-C**). Differential gene expression analysis revealed several hundreds of genes being strongly upregulated in *Leo1L* and *Paf1L* mutants; a notably higher number compared to bulk testis transcriptome analysis (**Figure 4E**). Furthermore, among the down-regulated genes, we found *kl-5* (**Figure 4E-F**). *kl-5* is one of six genes on the *D. melanogaster* Y chromosome classically known as male fertility factors ^44,45^, each essential for male fertility. In fact, all six male fertility factor genes showed reduced expression in *Leo1L* and *Paf1L* mutants in RNA-seq from spermatocyte as well as bulk testis, while *Ctr9L* mutants showed a similar trend with small effect size (**Figure 4G, Figure S14E**). All six fertility factor genes harbor gigantic introns in the megabase-range ^46,47^ that require the full ∼90 hours-duration of spermatocyte development to transcribe ^48^. We therefore speculate that tPAF, through an unknown co-transcriptional mechanism, enables expression of these gigantic genes on the Y chromosome, which likely contributes to the observed fertility defect in tPAF-deficient male flies. In sum, we find that tPAF harbors subunit-specific subfunctionalities related to nucleolar accumulation and genome regulation with especially Leo1L and Paf1L functioning to maintain cell type-specific gene expression in spermatocytes.

### tPAF ensures efficient transcription termination at testis-specific genes

Manual inspection of the genome-mapped RNA-seq coverage revealed a common pattern in which the upregulated genes (i) showed no expression in testes under wildtype conditions and (ii) were often positioned downstream of a highly expressed gene oriented in the same direction (**Figure 5A; Figure S17A-C**). We therefore hypothesized that read-through transcription from an upstream gene (the *read-through gene*) causes the upregulation of the downstream *read-in gene*. Of note, inspection of such individual loci revealed that while the increased expression is most pronounced in *Paf1L* and *Leo1L* mutants, *Ctr9L* and *Cdc73L* mutants show similar transcriptome phenotypes with lower effect size **(Figure 5A; Figure S17A-C)**. Consistent with a model of upregulation by read-through transcription from such upstream genes, a metagene analysis showed increased spermatocyte RNA-seq coverage immediately upstream of the upregulated *read-in genes* in *Paf1L* and *Leo1L* mutants compared to control libraries (**Figure 5B; Figure S17D, middle panels**). Furthermore, increased coverage in tPAF mutant conditions was also observed immediately downstream of the nearest expressed upstream gene on the same strand (**Figure 5B; Figure S17D, left panels**). We therefore conclude that read-through events from these upstream genes are the likely origins of the read-in transcription leading to the many upregulated genes in *Leo1L* and *Paf1L* mutants.

**FIGURE 5.**
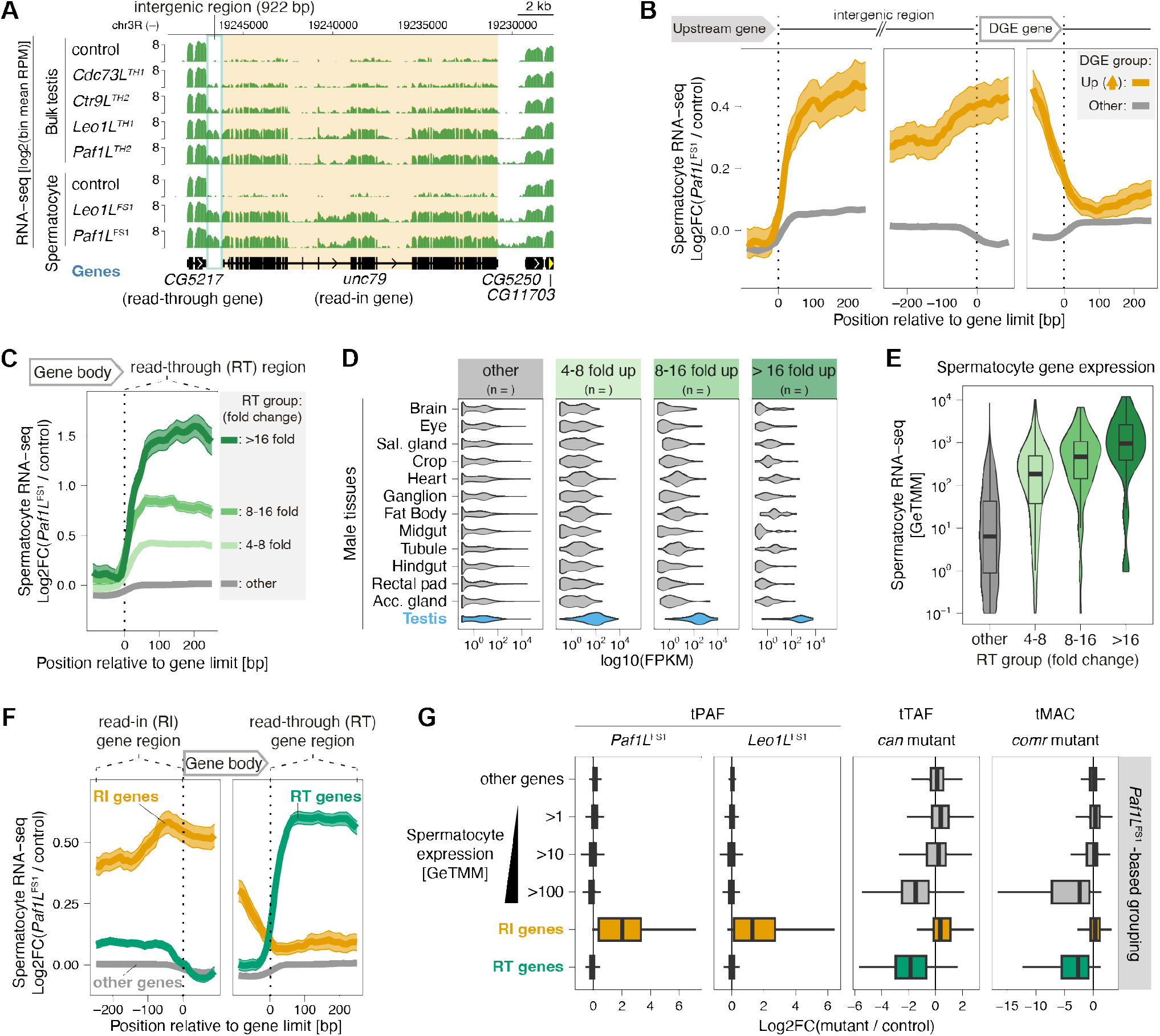
tPAF ensures efficient transcription termination at testis-specific genes. (A) Genome tracks showing RNA-seq coverage around the *unc79* gene for the indicated genotypes and sample type. The coverage signal is displayed as the log2 of mean reads per million mapped reads (RPM) per genome region bin. (B-C) Metagene analysis of RNAseq coverage fold change in Paf1L mutants relative to control samples. The standard error is plotted as a faded ribbon around the solid line showing the mean value at each genomic position. The plots compare coverage around bulk annotated genes (“other”) to that of genes upregulated in tPAF mutants (B) or to that of genes showing increased downstream RNA-seq signal in tPAF mutants (C). (D) Violin plot of gene expression levels based on FlyAtlas2 RNAseq from the listed fly tissues. Expression is plotted separately for the gene categories established in (C). FPKM = Fragments Per Kilobase per Million mapped fragments. (E) Violin plot of spermatocyte RNA-seq gene expression quantification of the gene categories established in (C). GeTMM = Gene length corrected trimmed mean of M-values. (F) Like (C-D) but here comparing read-in (RI) and read-through (RT) gene categories to the remaining (“other”) annotated gene loci. (G) Box plots showing the distribution of change in gene expression values between mutant and control RNA-seq samples for the listed gene categories and mutant genotypes. The data in the tPAF mutant panels is the spermatocyte RNA-seq data. The data underlying the tTAF and tMAC panels originates from reference ^49^.

Further inspection showed that the length of the intergenic space between the read-in gene, and the upstream read-through gene varies from a few hundred base pairs to more than 33 kilobases **(Figure S18A-H)**. Notably, for short (<1 kb) intergenic spaces, the RNA-seq coverage is similar across the intergenic space and exons of the *read-in gene* **(Figure S18A-D)**, while the RNAseq coverage at longer intergenic spaces is lower than the read-in exon coverage **(Figure S18E-H)**. This pattern was also found at loci where read-through from a *read-in gene* resulted in a secondary *read-in gene* activation **(Figure S18I-J)**. These results suggest that for short intergenic spaces, the spliced read-in gene mRNA may exist as a fusion transcript with the upstream mRNA, while for longer intergenic spaces most of the *read-in gene* mRNA likely has a presumably uncapped 5’ terminus close to the annotated transcription start site. Furthermore, RNAseq coverage over the *read-through genes* was unaffected by tPAF mutation and was orders of magnitude higher than over the downstream *read-in genes* **(Figure S18A-H)**, indicating that only a minor fraction of the transcription events at *read-through genes* are responsible for the observed *read-in gene* activation.

To characterize the genes that produce read-through transcription in tPAF mutants, we grouped annotated genes by the fold increase in RNA-seq coverage immediately downstream of the genes **(Figure 5C; Figure S19A)**. Genes with tPAF-dependent read-through show a strong tendency to be expressed specifically in the testis **(Figure 5D; Figure S19B)** and tend to be among the most highly expressed genes in spermatocytes **(Figure 5E)**. Since these patterns are observed for mild to strong read-through signature in both *Paf1L* and *Leo1L* mutants, we define read-through (RT) genes as spermatocyte-expressed genes having at least four-fold increase in downstream RNA-seq coverage in *Paf1L* mutants and read-in (RI) genes as genes at least four-fold increase in upstream RNA-seq coverage in *Paf1L* mutants and negligible expression in wildtype spermatocytes **(Figure 5F)**. Consistent with our observations from manual inspection, *read-in genes* are strongly upregulated in tPAF mutants while *read-through genes* are unaffected, as are other genes irrespective of their expression level in spermatocytes **(Figure 5G)**. Given the identified connection between tPAF and tTAF (**Figure 1A**), we next asked if read-through transcription in tPAF mutants occurs specifically at tTAF- or tMAC-regulated genes. Indeed, the vast majority of read-through genes also show strongly decreased mRNA expression in *can* (tTAF) mutants, as well as *comr* (tMAC) mutants compared to control (**Figure 5G**, right panels, data from ref. ^49^). tTAF/tMAC-dependent genes constitute many of the highest expressed genes in spermatocytes **(Figure 5G)**, which may explain why a dedicated transcription elongation complex is needed at these loci.

In sum, unlike canonical PAF1C, which promotes RNA Polymerase II elongation genome-wide^50^, tPAF enables efficient transcription termination specifically at tTAF/tMAC-dependent genes. While difficult to test directly, we speculate that the aberrant regulation of hundreds of genes caused by transcriptional read-through contributes to the sterility phenotype observed in *Leo1L* and *Paf1L* mutants through perturbation of the spermatocyte-type gene expression program.

### tPAF functionally connects transcription initiation factors and co-transcriptional effectors

The canonical PAF1 complex is thought to regulate multiple steps in gene expression by acting as a molecular landing pad for various effector proteins of co-transcriptional processing and chromatin regulation^32^. To uncover if the tPAF complex functions are enabled through a similar platform-like function, we performed proximity-labeling coupled to mass spectrometry analysis of the four tPAF-specific subunits, Paf1L, Leo1L, Ctr9L, and Cdc73L. Confocal imaging of testis from flies harboring both a GFP-nanobody–TurboID expression cassette and endogenously tagged tPAF genes showed a strong enrichment of biotinylation colocalizing with GFP-tagged tPAF protein (**Figure S20A**). Mass spectrometry analysis of biotinylated proteins enriched by streptavidin-based purification revealed high enrichment of all four tPAF paralog subunits **(Figure 6A-B; Figure S21A-C)**, thus indicating successful biotinylation of tPAF-associated proteins. We note that Ski8 was not observed in these analyses and speculate that this was due to technical reasons given the strong Ski8 enrichment in the more stringent biochemical tPAF complex purifications (**Figure 3D-F; Figure S9A-B**). In addition to the tPAF proteins, the mass spectrometry data also uncovered multiple subunits of the tTAF complex and the two testis-specific bromodomain proteins, tBRD-1, and tBRD-2, which we also observed to associate with tTAF in proximity labeling experiments (**Figure 1A**). These results strongly support a model where the genes regulated by tPAF are defined through the association of tPAF with tTAF and tBRD proteins.

**FIGURE 6.**
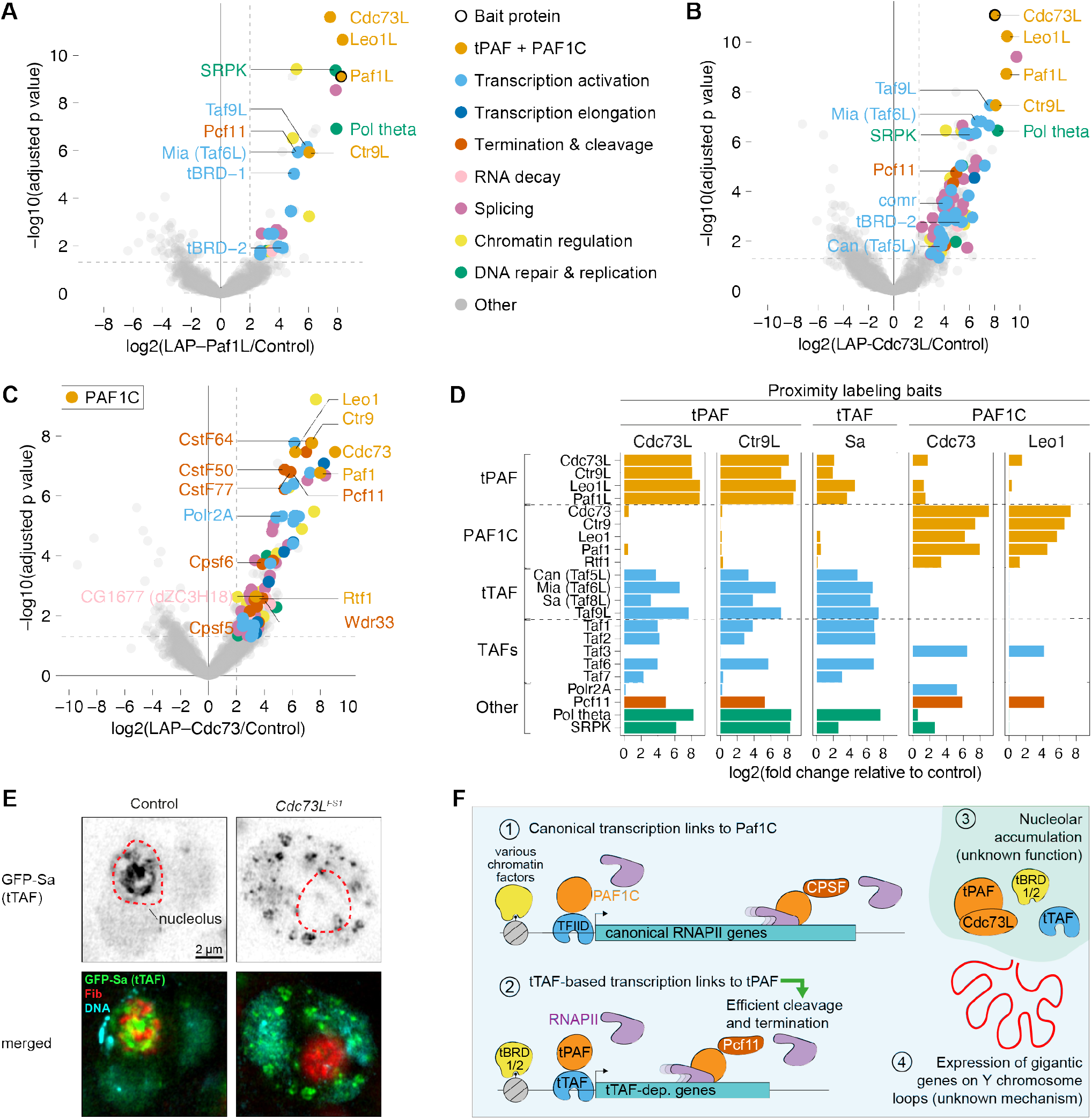
Protein associations support a multifunctional tPAF complex. (A-C) Enrichment values and corresponding statistical significance levels for proteins associated with the proximity labeling bait protein compared to control samples as detected by proximity labeling mass-spectrometry. The baits for the displayed data are Paf1L (A), Cdc73L (B), and Cdc73 (C). Colored highlights indicate manually annotated protein groups as listed to the right of (A). (D) Bar plot showing a comparison of enrichment for select proteins in different bait protein samples. Proximity labeling samples from tPAF, tTAF, and PAF1C bait proteins are compared as shown. (E) Confocal microscopy images showing the localization of GFP-tagged Sa (tTAF), Fibrillarin (Fib, anti-Fib immunoflourence staining), and chromatin visualized by DAPI. Control (*w*^*1118*^) and *Cdc73L* mutant genotypes are compared. Dashed red line: nucleolus border. (F) Schematic model comparing PAF1C and tPAF function. See discussion for details.

Notably, Pcf11, a key protein responsible for inducing eukaryotic RNA Polymerase II transcription termination^51,52^, was highly enriched by tPAF proximity labeling. This offers a mechanistic model of tPAF-mediated Pcf11 recruitment to mediate the observed tPAF function in transcription termination at tTAF-dependent genes. Supporting a multifunctional platform-like role of tPAF, we also observed the association of tPAF with proteins playing major roles in DNA maintenance (Pol Theta), chromatin regulation (NO66), and splicing (SRPK). Complementary characterization of canonical PAF1C protein associations by proximity labeling confirmed that tPAF and PAF1C do not share core subunits **(Figure 6C-D; Figure S21D, Figure S22)**. Furthermore, while some proteins, such as Pcf11, are associated with both tPAF and canonical PAF1C, comparative analysis uncovered that several proteins associate specifically with tPAF (e.g. tTAF, tBRDs, Pol Theta, and Taf2/6/7) or with PAF1C (e.g. Taf3 and all subunits of the Cleavage stimulation Factor (CstF) mRNA 3’ processing complex) **(Figure 6D; Figure S21E-F, Figure S22)**. Of note, only one of the thirteen RNA Polymerase II complex subunits (Polr2A) was enriched over control in only one set of samples (Cdc73, PAF1C) **(Figure 6C-D)**. This suggests that RNA Polymerase, similar to Ski8, may be difficult to detect due to technical reasons, or that most tPAF and PAF1C is mainly associated with the detected transcription factors and co-transcriptional effector proteins *in vivo*.

Further supporting a multifunctional role of tPAF, we also observed that tPAF, including Cdc73L, is required for the nucleolar localization of Sa-GFP (tTAF) protein **(Figure 6E; Figure S22B)**. *Cdc73L* mutants show hardly any defect in transcriptional regulation **(Figure S14D-E)** but are nonetheless fully sterile **(Figure 1E)**. Together, this indicates that tPAF, through Cdc73L-mediated nucleolar anchoring, serves a yet to be understood function in spermatocyte development.

In sum, we find that PAF1C and tPAF display distinct protein association profiles that provide a mechanistic model for the specialized functions in general RNA Polymerase II regulation and proper co-transcriptional regulation of spermatocyte-specific genes, respectively **(Figure 6F)**.

## DISCUSSION

### A paralogous Paf1 Complex facilitates male germline development

In this study, we identify tPAF, a paralog-based PAF1C-like complex, and characterize its essential function in germline development in *Drosophila* males. tPAF mutants displayed defects in germline development that indicate defects in meiosis during the spermatocyte stage **(Figure 1-2)**. Consistently, the tPAF proteins are expressed specifically in the pre-meiotic spermatocytes where they form a protein complex that additionally includes the Ski8 protein, which is also part of canonical PAF1C as well as the cytoplasmic SKI complex **(Figure 3)**. tPAF proteins are enriched in spermatocyte nucleoli but also accumulate in regions overlapping with chromosome territories. Likely independent of its putative function in nucleoli, we find that tPAF ensures efficient transcription termination at testis-specific genes that depend on tTAF for their transcriptional activation **(Figure 4-5)**. tPAF-induced transcription termination thereby prevents aberrant activation of hundreds of genes via read-in transcription. In addition, tPAF is required for expression of Y-chromosomal male fertility factor genes and thus altogether ensures cell type-specific gene expression in the male germline. Mechanistically, we propose that tPAF functions through protein associations that are conserved between tPAF and canonical PAF1C as well as tPAF-specific protein connections that have evolved after the duplication of the tPAF genes **(Figure 6)**.

### Cell type-specific regulation of transcription termination via tPAF

Canonical PAF1C functions to regulate RNA polymerase II transcription pause release, elongation, termination and 3’ processing and also affects local chromatin states (reviewed in ref. ^32^). On the contrary, we find that the tPAF complex has evolved to be a much more specialized regulator of gene expression as it (i) stimulates expression of the six gigantic fertility factor genes on the Y chromosome **(Figure 4E-G)** and (ii) facilitates efficient transcription termination specifically at tTAF-dependent genes **(Figure 5)**. We observed markedly stronger transcriptional defects in *Leo1L* and *Paf1L* mutants compared to mutants of the other tPAF components, *Cdc73L* and *Ctr9L*. This indicates that Leo1L and Paf1L proteins form a functional subcomplex, which is also supported by the close biochemical association of Leo1L and Paf1L proteins **(Figure 3G-I, Figure S9A)**. Notably, unlike the other canonical PAF1C proteins, human Leo1 and Paf1 have been found to accumulate at gene termini and knock-down of Leo1 and Paf1 elicits alternative polyadenylation defects in human cell culture^53^. Together with our findings, this indicates that regulation of RNA Polymerase II at gene termini is a conserved function of PAF1C, which in tPAF has evolved to enhance transcription termination at a specific subset of genes to maintain cell type-specific gene expression.

Through proximity labeling coupled to mass spectrometry analyses, we find that both tPAF and PAF1C complexes are associated with proteins of the cleavage and polyadenylation specificity factor (CPSF) complex **(Figure 6)**. However, while canonical PAF1C associates with various CPSF proteins, mainly Pcf11 is enriched in tPAF proximity labeling **(Figure 6A-C)**. Pcf11 is a central component of the CPSF complex but is known to play a major role in inducing transcription termination of RNA Polymerase II, even capable of inducing termination independent of the remaining CPSF complex proteins *in vitro*^*51,52*^. We speculate that the evolution of the association of specifically Pcf11 with tPAF compared to the CPSF-connected PAF1C reflects an adapted functional difference between the two paralogous complexes. Specifically, while tPAF depletion results in pronounced transcription termination defects, depletion of human Leo1L and Paf1L was reported to mainly cause alternative polyadenylation site usage^53^. The testis-specific genes at which tPAF ensures efficient termination are typically short and transcribed at very high levels **(Figure 5E)**. We propose that the very high transcription rates of spermatocyte-specific genes challenge cell type-specific gene regulation due to the risk of aberrant read-in gene activation, which tPAF counteracts through its ability to effectively induce transcription termination via Pcf11.

The mechanism underlying the observed requirement of tPAF for normal expression of the six male fertility factor genes remains unknown. However, we speculate that tPAF plays a role in the co-transcriptional regulation of these gigantic genes to facilitate their timely expression before the end of spermatocyte development^48,54^. For example, tPAF might enable timely RNA polymerase II transcription through the vast repetitive intronic sequences within the male fertility factor genes, or facilitate efficient splicing of these gigantic introns. Intron gigantism is also physiologically important in humans and other vertebrates^55^ and the *Drosophila* male fertility factor genes thus constitute a favorable model system for further investigations of the molecular mechanisms underlying expression of gigantic genes in general.

### Coupled initiation and termination in non-canonical transcription

While canonical PAF1C is thought to function at all RNA Polymerase II transcribed loci^53,56–58^, tPAF mutants display aberrant transcription specifically at tTAF-dependent genes **(Figure 6)**. Since tPAF is not required for the expression of tTAF-dependent genes **(Figure 5D-E)**, this argues that transcription activation via tTAF couples to tPAF-mediated transcription termination. Such coupling between a variant basal transcription factor complex and variant co-transcriptional processing factors has already been shown to enable important variations in RNA Polymerase II function. For example, proper snRNA 3’ processing relies on snRNA gene promoter elements, which are activated by the snRNA-activating protein complex (SNAPc)^59–61^, and in *Drosophila*, a TFIID variant complex couples heterochromatic piRNA precursor transcription to suppression of both splicing and polyadenylation site recognition as well as to a dedicated piRNA precursor nuclear export pathway^4,13,14,62,63^. tTAF-coupled transcription termination via tPAF thus supports a broader model for RNA Polymerase regulation, where variant transcription initiation mechanisms direct downstream processes in gene expression. Notably, alternative cleavage and polyadenylation is known to enable cell type-specific mRNA isoform expression^64^ and recent findings indicate that connections between transcription initiation and termination are widespread in the human genome^65^. The underlying molecular connection mediating these couplings, including possible tTAF-tPAF-like transcription and termination factor associations, remains to be uncovered. For tPAF, our proximity labeling results indeed support a model where recruitment to tTAF-dependent loci happens through direct protein-protein interactions with either tTAF or tBRD proteins **(Figure 1A and 6)**. Direct biochemical investigation of this model will be important to uncover how transcription initiation and termination processes are coupled.

*Cdc73L* and *Ctr9L* mutants display only mild read-through transcription phenotypes but still have meiotic defects and are sterile. Therefore, we currently cannot determine to which extent read-through transcription in the *Leo1L* and *Paf1L* mutants contributes to the sterility phenotype. Given that the read-through phenotype activates hundreds of genes aberrantly via read-in transcription **(Figure 5C)** the developmental impact of this effect is difficult to dissect experimentally but such a level of aberrant gene expression is in itself likely to disturb normal development. In addition, *Paf1L* mutants, which perturb both tPAF integrity and transcription termination **(Figure 4, 5, S15)** show overall stronger developmental phenotypes compared to *Cdc73L* mutants **(Figure 2, S4, S6)** which have only very mild transcription termination defects **(Figure 4E, 5B)**. Therefore, we propose that Leo1L and Paf1L function together to ensure cell identity maintenance by blocking promiscuous gene activation that would otherwise arise through read-in transcription from upstream tTAF-dependent genes.

### Open questions and future directions

Maintenance of cell type-specific gene expression through repression of lineage-incompatible genes is also known from the Kumgung (Kmg) zinc finger protein acting through the dMi-2 repressor in the *Drosophila* male germline^66^ and from the separate CoREST complexes functioning in *Drosophila* somatic and germline cells^67^. The combined action of transcription activation and repression mechanisms thus seems particularly important for germ cell maintenance, begging future investigation of similar pathways beyond *Drosophila*. We propose that paralog-based gene regulatory complexes form connected functional pathways and that such connections are required to fix the problems arising from bending or bypassing the textbook dogmas of gene regulation. Such paralog-based pathways represent valuable model systems for future mechanistic studies of animal germline genome regulation as well as of the parental canonical complexes through comparative mechanistic studies.

## ACKNOWLEDGEMENTS

We thank P. Duchek and J. Gokcezade (IMBA fly facility) for generating CRISPR-edited tPAF frame-shift mutant flies during Peter Andersens’ affiliation with IMBA. We thank Changwei Yu (Brennecke group, IMBA) for sharing the TurboID-GFPvhh fly line ahead of publication. We thank Mie Aarup (Andersen group, MBG-AU) for organizing essential lab infrastructure for the project and for contributing help with plasmid cloning. We much appreciate the helpful comments on the manuscript from Torben H. Jensen and Anne F. Nielsen. Much of the computing for this project was performed on the GenomeDK cluster and we thank GenomeDK and Aarhus University for this support. Flow cytometry and cell sorting were performed at the FACS Core Facility, Aarhus University, Denmark. In the Andersen group, this work was supported by the Novo Nordisk Foundation (NNF18OC0030954), an AIAS Marie Curie COFUND Fellowship, and the Independent Research Foundation Denmark (9064-00056B). The Hayashi group is supported by the Australian Research Council (DP210102385).

## AUTHOR CONTRIBUTIONS

A.P.V. performed the experimental work apart from the contributions listed below, analyzed data, and drafted the manuscript.

A.G. carried out the characterization of tPAF mutant developmental and meiotic phenotypes.

A.J.R. performed all experiments related to proximity labelling and Ski8.

A.E. carried out tissue preparation and fluorescence-activated cell sorting (FACS) for the generation of spermatocyte RNA-seq

S.R. made the structure modelling of tPAF and PAF1C complexes using Alphafold2.

M.V.L. carried out the mass spectrometry data acquisition.

R.H. conceived the study, performed data analyses, wrote the manuscript, and supervised A.G.

P.A. conceived the study, performed data analyses, wrote the manuscript, and supervised A.P. V., A.J.R., A.E, and S.R.

## MATERIALS AND METHODS

### Data and code availability

The high-throughput sequencing data generated for this publication has been deposited at the Sequence Read Archive (SRA) under accession number: GSE263955

The mass spectrometry proteomics data have been deposited to the ProteomeXchange Consortium via the PRIDE partner repository with the dataset identifiers PXD049360 (co-IP data) and PXD049375 (proximity labeling data). Interactive volcano plots can be accessed from the GitLab below project and through this webpage: http://genome-ftp.mbg.au.dk/public/PAN/Data_tPAF_Vilstrup_2024/Interactive_mass_spec_plots/

Computational scripts used for analysis and data visualization, processed data tables used for plotting, and interactive visualization of the mass spectrometry data can be accessed here: https://gitlab.com/AndersenGermlineLab/tpaf_vilstrup_2024

### Drosophila maintenance and strains

*D. melanogaster* strain maintenance, as well as experiments, were done on standard medium at 25°C at 70% humidity with 12:12 hrs light/dark cycle unless otherwise stated. Fly strains used in this study are listed in Supplemental Table 1. Frame-shift mutants of tPAF genes were generated by Peter Duchek, Vienna Biocenter Core Facility. Knock-in (KI) D. melanogaster lines tagging the endogenous gene tPAF and PAF1C genes with N-terminal 3xFLAG-3xV5-GFP were generated by Rainbow Transgenic Flies, Inc, CRISPR cloning. Eric Wieschaus kindly gifted the RFP-tagged Fib fly strain. GFP-tagged Tpl94D and ProtA strains were kindly gifted by Benjamin Loppin, University of Lyon. tMAC and tTAF mutants were kindly gifted by Margaret T. Fuller, Stanford University. We used FlyBase^68^ to find information on phenotypes/function/stocks/gene expression (etc) of genes mentioned in this study.

### Paralog identification

Paralog genes of PAF1C and ortholog genes across species were identified by NCBI tBLASTn using the RefSeq genome database with default search parameters^69^. Potential orthologs were verified with BLASTp search within the *Drosophila melanogaster* genome to verify that the correct ortholog was the top hit. Orthologs were further verified by comparing the synteny of the adjacent genomic region. Genes were accepted as orthologs if they were the top hit in BLAST searches and had comparable synteny. Gene identifiers of paralogs and orthologs are listed in Supplementary Table 2.

### Fertility assay

Fertility assays were done with transheterozygous mutant males or females mated to control (*w*^*1118*^) flies of the opposite sex. Matings were set up with flies of 2-4 days old in a 4:8 ratio of males to females in a food vial for 24 hrs. After 24 hrs. flies were transferred to a petri dish with solidified agar and apple juice with yeast paste and net covering. Flies were allowed to lay eggs on the agar for 24-30 hrs., whereafter they were removed, the petri dish was covered and left for a further 24 hours to let larvae hatch from the eggs. Hereafter, the ratio of hatched vs. unhatched out of 100 eggs was counted. Each tPAF mutant was tested with two different transheterozygous mutant genotypes and the fertility of each genotype was measured in duplicates.

### Immunostaining of post-meiotic spermatids

The immunostaining protocol was adapted from reference ^70^. Briefly, a testis pair from young tPAF1C mutants and control (*w*^*1118*^) flies was dissected and placed on a glass slide with 20 μL of 1xPBS. The posterior end of the testis was nicked using a needle to release elongated spermatids. A coverslip was gently placed atop, and excess PBS was removed using Kimwipe to obtain a monolayer squash. The samples were quickly snap-frozen in liquid nitrogen for 30 seconds. The coverslip was immediately removed, and samples were immersed in 95% ice-cold ethanol, incubating for 20 minutes at -20°C. The samples were then fixed using 4% formaldehyde in 1x PBS-0.3% Triton X-100 for 7 minutes at room temperature. After two washes with 1x PBS for 5 minutes each, the samples were permeabilized in 1x PBS-0.3% v/v Triton X-100 for 30 minutes at room temperature. Subsequently, the samples were washed with 1x PBS twice for 5 minutes and incubated with DAPI (1 mg/ml, 1:100) and Alexa Fluor 488 phalloidin (1:1000, Thermo Scientific, A12379) for 30 minutes at room temperature. Finally, samples were washed with 1x PBS and mounted using VECTASHIELD antifade mounting medium (VectorLabs, H-1000-10).

For imaging the polarization process, five pairs of testis tissue from young tPAF1C mutants and control *w*^*1118*^ flies were dissected and transferred into an Eppendorf tube containing fixation solution (2% formaldehyde in 1X PBS 0.3% Triton X-100) for 10 minutes at room temperature. The samples were washed thrice with 1x PBS-0.3% Triton X-100, each for 10 minutes. Subsequently, the samples were permeabilized in 1x PBS-1% Triton X-100 for 30 minutes, followed by two washes using 1x PBS-0.3% TritonX, each for 10 minutes. The samples were then blocked with 1x PBS-0.3% Triton X-100 containing 0.05% w/v BSA for 30 minutes at room temperature. Samples were incubated with the primary antibody Orb 4H8 (mouse) from DHSB (1:1000) in 1x PBS-0.3% Triton X-100 containing 0.05% BSA overnight at 4°C. After three washes with 1x PBS-0.3% Triton X-100, each for 10 minutes, the samples were incubated with DAPI (f.c. 10 μg/mL) and anti-mouse IgG, Alexa Fluor® 647 Conjugate (1:1000) for 30 minutes at room temperature. Finally, samples were washed with 1x PBS and mounted using VECTASHIELD antifade mounting medium (VectorLabs, H-1000-10).

For imaging the chromosomal condensation process, five pairs of testis tissue from 0-2 day old tPAF1C mutants and *w*^*1118*^ with His2AV-RFP and beta-Tubulin-GFP were dissected and transferred into an Eppendorf tube containing fixation solution (2% formaldehyde in 1X PBS 0.3% Triton X-100) for 10 minutes at room temperature. The samples were washed thrice with 1x PBS-0.3% Triton X-100, each for 10 minutes, and mounted using VECTASHIELD antifade mounting medium (VectorLabs, H-1000-10). All samples were imaged using a Zeiss LSM 800 laser scanning confocal microscope, and images were processed in Fiji^71^. Pro-metaphase I and metaphase I identification were based on the formation of astral fibers around the nucleus and the spindle formation, respectively.

### Live imaging

Live imaging was performed to examine whole testis tissue, nebenkern stage, and meiosis I prometaphase. Testes tissues from 0-2 day old male flies were dissected from the described genotypes and placed on a glass slide with 20 μL of 1x PBS. For imaging the nebenkern stage and meiosis I, the anterior end of the testis was nicked using a needle to release the cells from the dissected tissue, while the tissue was left intact for whole testis overviews. A coverslip was gently placed on the dissected tissue, and excess PBS was removed using a Kimwipe to obtain a monolayer squash. The samples were imaged via an Olympus IX51 phase contrast microscope within 10 minutes of slide preparation. Images were processed in Fiji^71^.

### Fixed immunofluorescence sample preparation and imaging

Testis were dissected from males of age 1-3 days in cold 1xPBS on ice. Tissues were fixed in 2% formaldehyde solution for 20 minutes at room temperature with nutation. After fixation tissues were washed 3x10 min in PBX (1xPBS with 0.3% Triton X-100), blocked in BBX (PBX with 1% BSA) for 30 min, and washed once in PBX for 10 min, all steps with nutation at room temperature. Tissues from Knock-in LAP-tagged PAF1C flies were detected with anti-V5 tag (Invitrogen, R96025) 1:500 in BBX incubated overnight at 4°C with nutation. Hereafter tissues were washed 4x10 min in PBX with nutation at room temperature. Samples were then incubated with anti-mouse IgG, Alexa Fluor® 647 Conjugate (Cell signaling, 4410S) 1:1000 in BBX overnight at 4°C with nutation. Tissues were washed for 10 min in PBX, 5 min with 1:10 000 DAPI, and 2x10 min in PBX at room temperature with nutation before being mounted on a microscopy slide with ProLong Diamond mounting medium (Thermo Fisher Scientific, P36961). All other tissues were not incubated with antibodies, only stained with DAPI following the steps described for KI-PAF1C samples. Samples were imaged using Zeiss LSM800 confocal microscope with airyscan (Bioimaging Core Facility at Aarhus University, https://imaging.au.dk). Images were processed in Fiji^71^.

### Whole fly sample preparation for co-IP

Knock-in LAP-tagged tPAF flies and control flies expressing the LAP tag only (“N1NLS” stock) were collected when 1-3 days old and snap-frozen in liquid nitrogen. Samples were made into powder by grinding at 400 rpm for three times one minute in a Planetary Ball Mill PM 100 machine (Retsch). To avoid degradation during milling, the samples were cooled by liquid nitrogen through sample handling and processing. For co-IPs, each powdered sample was dissolved in five powder sample volumes of protein lysis buffer (20 mM Tris-HCl pH 7.5, 150 mM NaCl, 2 mM MgCl2, 10 % Glycerol, 1 mM DTT, 1 mM PefaBloc SC Plus, 0.2 % IGEPAL CA-630, ELGA water). Samples were then lysed by incubation on ice for 15 minutes, centrifuged for 5 minutes at 4°C, and lysate supernatant was transferred to clean tubes. 8 μL anti-FLAG M2 magnetic beads (Sigma-Aldrich, M8823) were added to each sample and the lysate/antibody mixtures were incubated with rotation for three hours at 4°C. The magnetic bead-coupled antibodies were then washed four times two minutes in 1 mL protein lysis buffer followed by five brief washes with 1 mL wash buffer (20 mM HEPES pH 7.4, 2 mM MgCl2, 150 mM NaCl, ultrapure water (ELGA)). In between washes, the beads were quickly spun down and pelleted on a magnetic rack to remove supernatant. Lastly, the beads were pelleted, the supernatant removed, and the samples were snap-freezed in liquid nitrogen and stored at -70°C until mass spectrometry analysis.

### Biotin proximity labeling sample preparation

1-3-day-old male flies harboring knock-in LAP-tagged tPAF genes combined with a transgenic biotin ligase fused to a GFP-nanobody (TurboID-GFPvhh)^31^, were transferred from standard cornmeal food to biotin-supplemented food (100 μM) for 24 hours before testis dissection. 200 testis pairs were dissected in ice-cold 1xPBS. PBS was exchanged with 100 μL lysis buffer (50 mM Tris-HCL pH 7,5, 150 mM NaCl, 0,1 % SDS, 1 % Triton X-100, 0,5 % Na-deoxycholate, Benzonase, 1X cOmplete, and 1mM DTT) and the tissue was homogenized using an electric pestle on ice for 20 seconds followed by quick centrifugation at 20.000 g at 4°C. The pestle treatment was repeated. The lysates were incubated at 4°C for 20 minutes followed by centrifugation for 10 minutes at 20.000g at 4°C. 150μL of Pierce™ magnetic streptavidin beads (S88817) pre-equilibrated in lysis buffer were mixed with the lysates and incubated overnight at 4°C with nutation. The beads were washed consecutively under rotation for 10 minutes in 1 mL of each of the following buffers at 4°C: (i) Lysis Buffer, (ii) 2% SDS in Tris-EDTA pH 7,5 (wash done at room temperature to avoid SDS precipitation), (iii) Wash Buffer 1 (50 mM HEPES pH 7,5, 500 mM NaCl, 1 mM EDTA, 0,1% Na-deoxycholate, 1% Triton X-100, and Benzonase), (iv) Wash Buffer 2 (10 mM Tris-HCl pH 7,5, 250 mM LiCl, 1 mM EDTA, 0,5 % Na-deoxycholate, and 1 % NP40) and (v) lastly six times with Wash Buffer 3 (20 mM Tris-HCl pH 7,5 and 137 mM NaCl). The beads were frozen in liquid nitrogen and shipped on dry ice for mass spectrometry analysis.

### Sample preparation for mass spectrometry

Mass spectrometry was done at DTU Proteomics Core, Kgs. Lyngby, Denmark. The samples were mixed with 20 μL lysis buffer (6 M Guanidinium Hydrochloride, 10 mM TCEP, 40 mM CAA, 50 mM HEPES pH 8.5) and sonicated on high for five times 30 seconds in a Bioruptor sonication water bath (Diagenode) at 4°C. Samples were diluted 1:3 with 10% Acetonitrile, 50 mM HEPES pH 8.5, and 200 ng LysC (MS grade, Wako) was added (1:50 enzyme to protein ratio). The samples were then incubated at 37°C for four hours. The samples were further diluted 1:10 with 10% Acetonitrile, 50 mM HEPES pH 8.5, and 100 ng trypsin (MS grade, Promega) was added (1:100 enzyme to protein ratio) followed by incubation overnight at 37°C. Enzyme activity was quenched by adding 2% trifluoroacetic acid (TFA) to a final concentration of 1%. The samples were desalted on SOLAμ SPE plate (HRP, Thermo), following the standard procedure. The dried peptides were reconstituted in 12 μL 2% ACN, 1%TFA/20 μL HEPES 50 mM pH 8.5.

### Mass Spectrometry Analysis

Peptides were loaded onto a 2 cm C18 trap column (ThermoFisher 164946), connected in-line to a 15 cm C18 reverse-phase analytical column (Thermo EasySpray ES904) using 100% Buffer A (0.1% Formic acid in water) at 750 bar, using the Thermo EasyLC 1200 HPLC system, and the column oven operating at 35°C. Peptides were eluted over a 140-minute gradient ranging from 6 to 60% of Buffer B (80% acetonitrile, 0.1% formic acid) at 250 nL/min, and the Q-Exactive instrument (Thermo Fisher Scientific) was run in a DD-MS2 top10 method. Full MS spectra were collected at a resolution of 70,000, with an AGC target of 3×106 or a maximum injection time of 20 ms and a scan range of 300–1750 m/z. The MS2 spectra were obtained at a resolution of 17,500, with an AGC target value of 1e6 or maximum injection time of 60 ms, a normalized collision energy of 25, and an intensity threshold of 1.7e4. Dynamic exclusion was set to 60 s, and ions with a charge state <2 or unknown were excluded. MS performance was verified for consistency by running complex cell lysate quality control standards, and chromatography was monitored to check for reproducibility.

### Analysis of mass spectrometry data

Mass spectrometry data were analyzed using the FragPipe (https://github.com/Nesvilab/FragPipe) proteomics pipeline using standard parameters for label-free quantification unless otherwise noted. Fully detailed search and analysis parameters are available through the listed PRIDE repository accessions. Briefly, the Uniprot Drosophila melanogaster proteome database was added 50 % reversed decoys, and spectra were searched against the proteome database using MSFragger^72^ with a wide precursor mass tolerance (+/- 50 PPM). Methionine oxidation and N-terminal acetylation were set as variable modifications and a maximum of two missed trypsin cleavages was allowed. Peptide-to-spectrum matches (PSM) were validated and rescored using Percolator^73^ and MSBooster^74^, respectively, with standard settings. Protein inference was done using ProteinProphet^75^. Label-free quantification was performed using IonQuant^76^ with match-between-runs (MBR).

Statistical analysis of the FragPipe search and quantification results were performed using FragPipe-Analyst (https://github.com/MonashProteomics/FragPipe-Analyst) with the following settings: Datatype = LFQ, Intensity type = Intensity, Normalization type = no normalization, Imputation type = minimum (co-IP data) or Perseus-type (proximity labeling data), Type of FDR correction = Benjamini-Hochberg. The applied parameters were chosen based on extensive parameter testing and evaluation of signal-to-noise ratios based on expected intra-complex protein-protein interactions.

### AlphaFold prediction

Homology model of D. melanogaster tPAF complex comprising the proteins Leo1L, Paf1L, Ctr9L, Cdc73L and Ski8 and PAF1C complex comprising Cdc73, Ctr9, Leo1, Paf1, and Rtf1 were manually assembled based on the cryoEM structure of human PAF complex^77^. 3D coordinates from individual *D. melanogaster* proteins were extracted from the AlphaFold database^78^ and the main domains were superimposed onto the respective human PAF counterparts using PyMOL v2.5. For Leo1L and Paf1L, manual adjustments were made using Coot^79^ to fit the unstructured regions to resemble interactions found in the human PAF complex.

### Sample preparation for whole-testis RNA-seq library

RNA was isolated from testis tissues of 1-3 days old transheterozygous tPAF mutants and control (*w*^*1118*^) flies. Tissues were dissected in cold 1x PBS on ice. Samples were homogenized and lysed in 1mL TRIzol. 200 μL chloroform was added, samples were vigorously shaken, incubated at room temperature for 3 minutes, and centrifuged for 15 minutes at 12 000g at 4°C. The RNA-containing phase was transferred to a clean RNase-free tube with 1 μL GlycoBlue co-precipitant (Thermo Fisher, AM9516), 550 μL isopropanol was added and samples were incubated overnight at -20°C. The following day samples were centrifuged at 16 000g for 30 minutes at 4°C and supernatant was removed from the RNA pellet. 1 mL ice-cold 75% ethanol was added to the RNA pellet, samples were centrifuged at full speed for 5 minutes at 4°C, and the supernatant was removed. Samples were briefly centrifuged at full speed at 4°C and residual ethanol was removed from the RNA pellet. Samples were dissolved in 15 μL RNAse free water. Genomic DNA was removed from the purified RNA samples using RNA Clean & Concentrator™-5 columns with on-column DNase digest (Zymo Research, R1014). For rRNA-depleted samples, rRNA was removed with RNase H using a modified version of the original protocol in ^80^ as described in ^13^. Briefly, rRNA depletion was performed using a mixture of antisense oligos complementary to the mature Drosophila melanogaster rRNA transcripts. The oligo pool was mixed with 1 μg DNAse-free total RNA in RNase H Buffer (20 mM Tris-HCl pH 8, 100 mM NaCl) and annealed over a temperature gradient from 95°C to 45°C. The RNA-DNA hybrids were then digested with thermostable RNase H (NEB, M0523S) at 45°C for 1 hour. Next, RNA was purified and the DNA oligos were removed from samples using RNA Clean & Concentrator™-5 columns with on-column DNase digest (Zymo Research, R1014) according to the manufacturer’s instructions. rRNA depletion was verified by qPCR. Library preparation was done with the NEBNext Ultra II Directional RNA Library Prep Kit for Illumina (NEB, E7770L), following the recommended kit protocol. Amplified libraries were purified by SPRI beads and library quality was assessed with Bioanalyzer High Sensitivity DNA Analysis (Agilent Technologies, 5067-4626). Samples were sequenced on a NovaSeq 6000 sequencing system (Illumina).

### Sorted spermatocyte RNA-seq library preparation

For spermatocyte isolation, testis tissues were dissected from 1-3 days old knock-in LAP-Cdc73L flies (control or in combination with *Paf1L* or *Leo1L* mutant alleles) into cold Schneider’s medium on ice. Following, the medium was removed and the tissue was homogenized with a plastic pestle and lysis buffer (560 μL PBS, 70 μL 1x trypsin, 70 μL 2% collagenase) was added. Samples were rotated at room temperature for 30 minutes followed by careful pipetting with a p1000 tip. The cell suspension was then drawn through a 40 μm mesh cell strainer into 500 μL Schneider’s medium. 100 μL additional Schneider’s medium was drawn through the cell strainer to collect the remaining cells. Samples were centrifuged at 1200 rpm for 7 minutes at 4°C and the supernatant was removed. Then, the cells were resuspended in 200 μL Schneider’s medium, and DNA was stained with bisbenzimide Hoechst 33342 (Sigma). GFP-expressing cells were then isolated by fluorescence-activated cell sorting (FACS) on a FACSAria III machine (BD Biosciences, FACS Core Facility at Aarhus University, www.facs.au.dk). Flowjo software (Treestar) was used for data analysis. The preparation and FACS sorting of control and mutant spermatocytes were performed in three independent experiments. To this end, we isolated control and mutant spermatocytes with a purity of at least 85% by FACS sorting and characterized their transcriptome by RNA-seq (see also **Figure S16**). Wildtype testes were used to clearly identify the GFP-expressing spermatocyte population during flow cytometry. RNA isolation was performed as described above for whole testis RNA-seq. Sorted spermatocyte RNA-seq libraries were prepared for each sample with and without rRNA depletion to minimize technical biases from working with small sample amounts. For the analyses, both library variants were pooled and analyzed together (see also the section on Analysis of RNA-seq data). Otherwise, the library preparation was performed as described above for whole testis RNA-seq.

### Analysis of RNA-seq data

After trimming to bases 5-45, RNA-Seq reads were mapped to the *Drosophila melanogaster* genome (dm6, r6.40) using STAR^81^ (for genome-mapping) or to *Drosophila melanogaster* mRNA sequences using Salmon^82^ (for transcriptome quasi-mapping). Normalization between samples was done based on the number of genome-unique mapping reads for each sample (for STAR-based mapping). For gene expression quantification, Gene length corrected TMM (GeTMM)^83^ values were calculated using edgeR^84^ (v3.34.0). Differential Gene Expression analysis was performed on Salmon analysis output using DESeq2^85^. The scripts used for RNAseq analysis and visualization as well as related processed data files are available at GitLab (see the ‘Data and code availability’ section above)

### Data visualization

All data panels, excluding schematics and microscopy images, were produced using R^86^, RStudio^87^, the seqNdisplayR package (genome coverage plots) ^88^, and the Tidyverse package assembly^89^.

