## Supplemental Figures for "A germline PAF1 paralog complex ensures cell type-specific gene expression"

### Figure S1

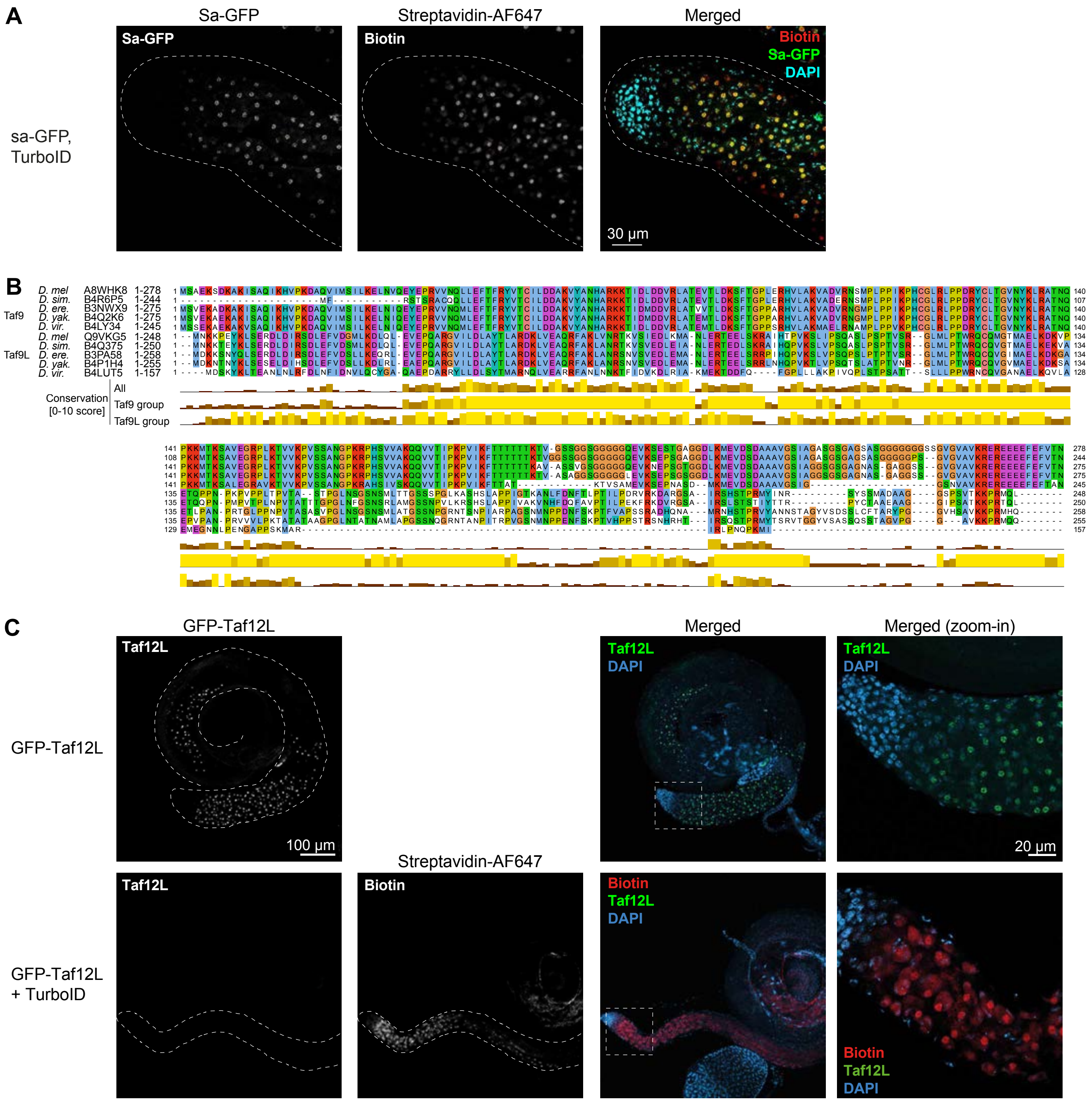


**FIGURE S1**

(A) Confocal microscopy images showing the apical tip of a testis from a Sa-GFP, TurboID fly. The images display localization of GFP-tagged Spermatocyte arrest (Sa; tTAF) protein, biotin accumulation (Streptavidin-AlexoFluor647), and chromatin visualized by DNA DAPI staining. The dashed line indicates the border of the testis tissue. (B) Protein sequence alignment of TAF9 and the identified paralog, CG14930 / TAF9L, from the indicated Drosophila species. The amino acid color code ("clustal") indicates similar amino acid properties. The conservation score below the alignment shows the percent divergence for the listed protein sequence groups. (C) Confocal microscopy images showing the whole testis tissue from flies harboring Taf12-GFP alone (top row) or combined with a TurboID cassette (bottom row). The images display localization of GFP-tagged Taf12L (tTAF) protein, biotin accumulation (Streptavidin-AlexoFluor647), and chromatin visualized by DNA DAPI staining. The dashed line indicates the border of the testis tissue. The rightmost images show zoomed-in merged-channel images of the testis apical tip.

### Figure S2

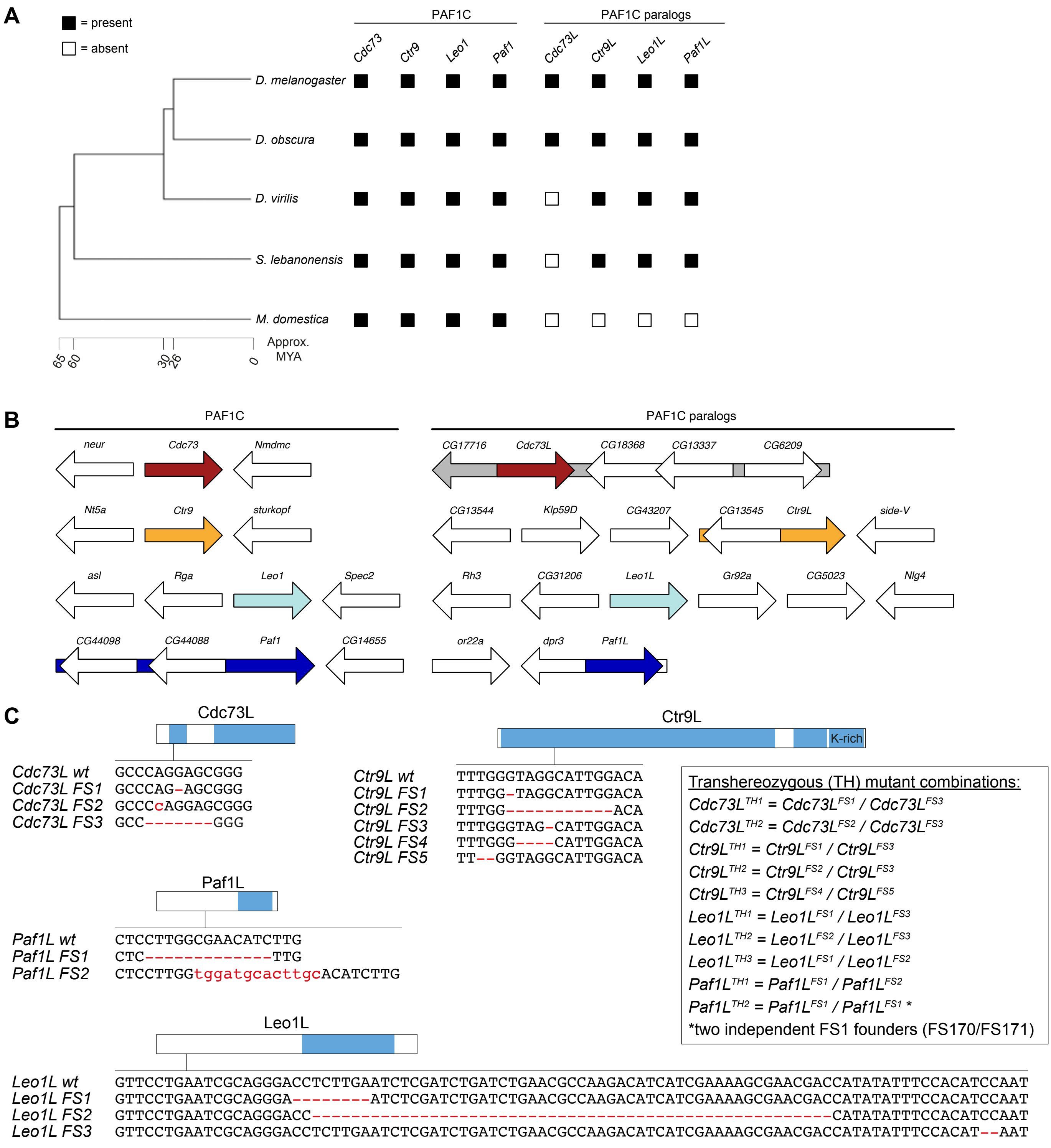


**FIGURE S2**
(A) Phylogenetic tree of the indicated species (left) with schematic indication of the presence (solid black box) or absence (white box) of PAF1C and tPAF genes as assessed by BLAST search and synteny analysis. Approx. MYA = approximated million years ago since the divergence of the phylogenetic tree branch. (B) Schematic representation of the gene annotation context of PAF1C and tPAF genes used for synteny analysis. (C) Schematic showing of tPAF genes with protein domains from Figure 1C indicated in blue. The nucleotide sequence changes in the CRISPR/Cas9-generated tPAF mutant alleles are illustrated below the gene schematics.

#### Figure S3

**A**

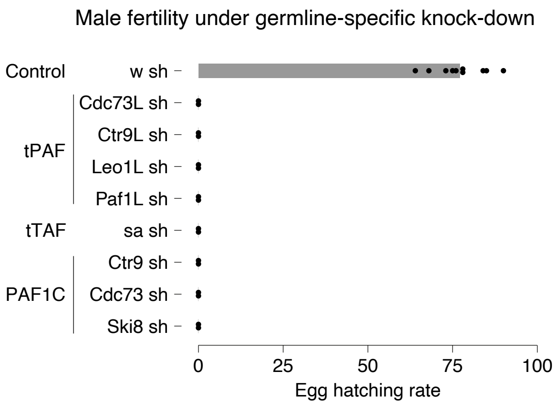


**FIGURE S3**
(A) Bar plot showing the mean hatching rate from crosses between control females and males subjected to either control or tPAF germline knock-down using bam>Gal4 driver combined with short hairpin (sh) RNA expression cassettes. Individual data points from biological replicates (n = 2) are shown as black dots.

#### Figure S4

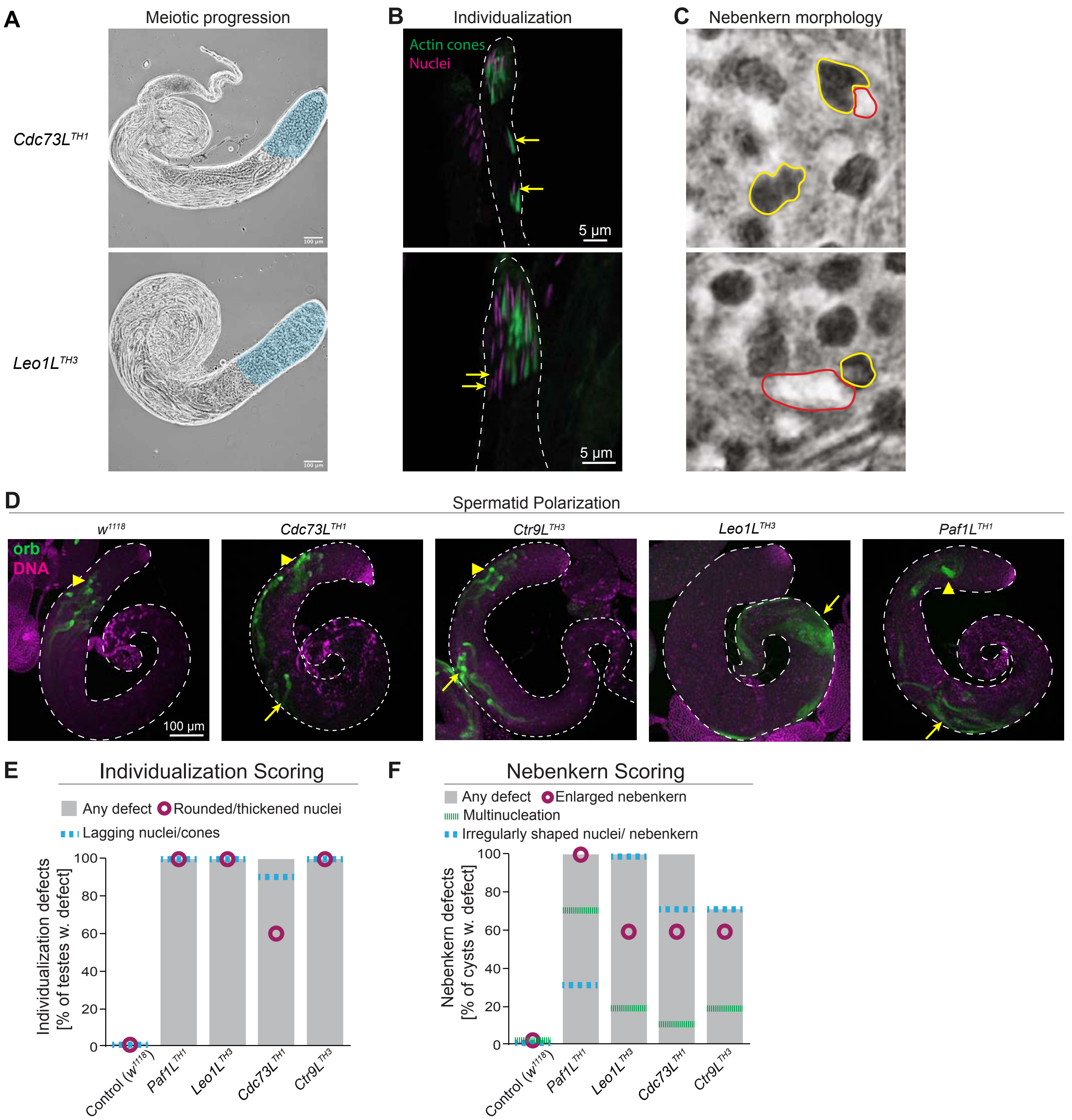


**FIGURE S4**
(A-C) Images of tPAF mutant testes (*Cdc73L^TH1^* and *Leo1L^TH3^*) showing (A) meiotic progression by phase contrast with pre-meiotic regions marked by cyan, (B) immunostaining images of elongating spermatids that are undergoing individualisation with lagging nuclei indicated by arrows, and (C) phase contrast images of nebenkern stage spermatids, showing irregularly shaped nuclei (red) and nebenkern (yellow). (D) Wild-type (*$w^{1118}$*) and the indicated tPAF mutant whole testes stained with anti-Orb antibodies, showing the mis-localisation of elongating spermatids. Wild-type elongating spermatids migrate to the apical end of the testis and the strong Orb staining is seen at the very apical end (arrowheads). In tPAF mutants, some elongating spermatids fail to reach the apical end, hence the strong Orb signals are seen in the middle part of the testis (arrows). (E-F) Bar plots showing the percentage of scored defects in spermatid individualization (E) and nebenkern/onion stage morphology (F) (n=10, representative of independent experiments). Gray bars indicate the frequency of any defect scored. Colored lines and circles show frequencies of individual defects (see top legend in each panel).

#### Figure S5

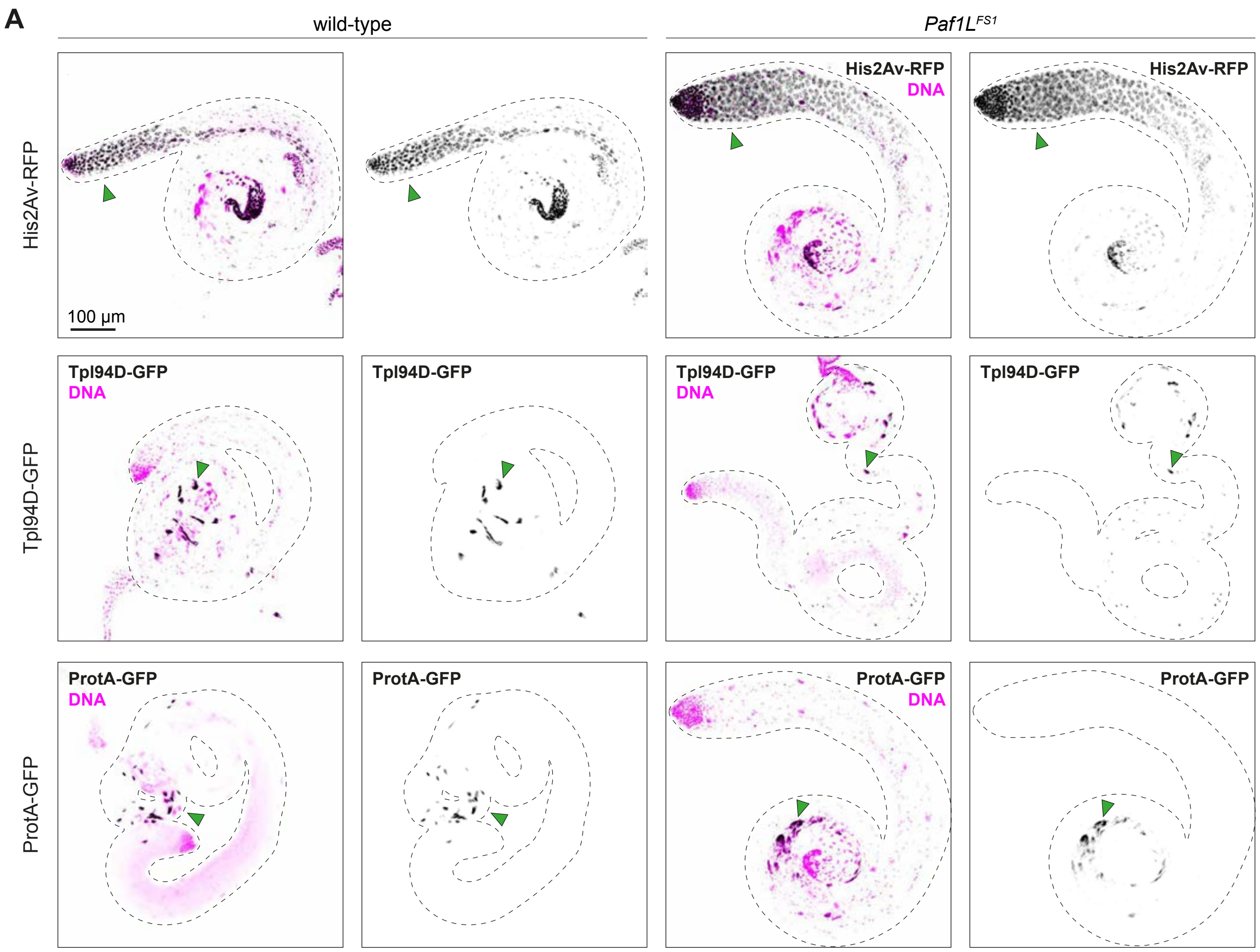


**FIGURE S5**
(A) Confocal images showing chromatin visualized by H2Av-RFP tagged histone proteins (top row), the transition protein Tpl94D-GFP (middle row), and protamine A (ProtA-GFP; bottom row) with
with DAPI-stained DNA shown in magenta. The leftmost images show testis from control flies, while the rightmost images show testis from Paf1L (tPAF) mutants.

### Figure S6

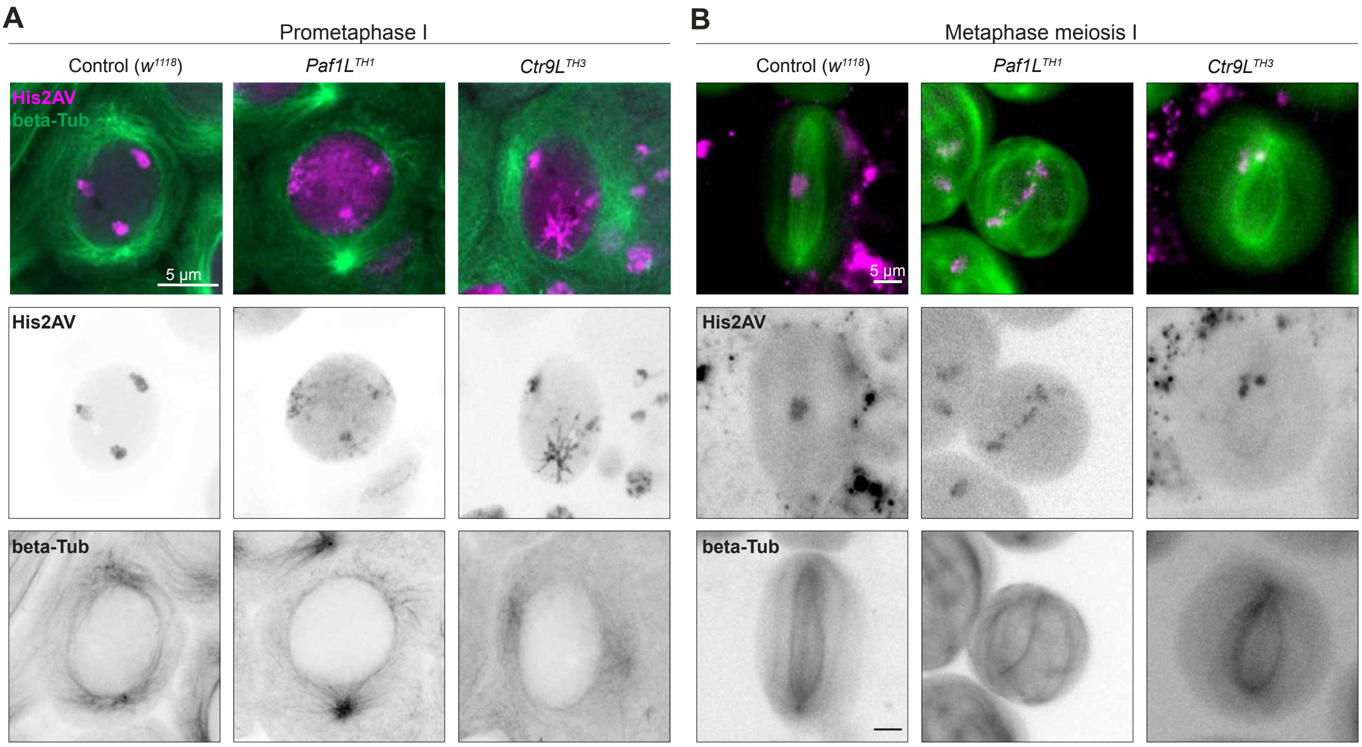


**FIGURE S6**
Individual colour channels of the images used in Figure 2G and H. Live (A) or fixed (B) cell imaging of spermatocytes at pro-metaphase I (A) and metaphase I (B) obtained through fluorescent microscopy, comparing wild type to tPAF mutants. Cells express RFP-tagged Histone H2Av and GFP-tagged beta-Tubulin to aid visualisation of the chromosomes and microtubules.

#### Figure S7

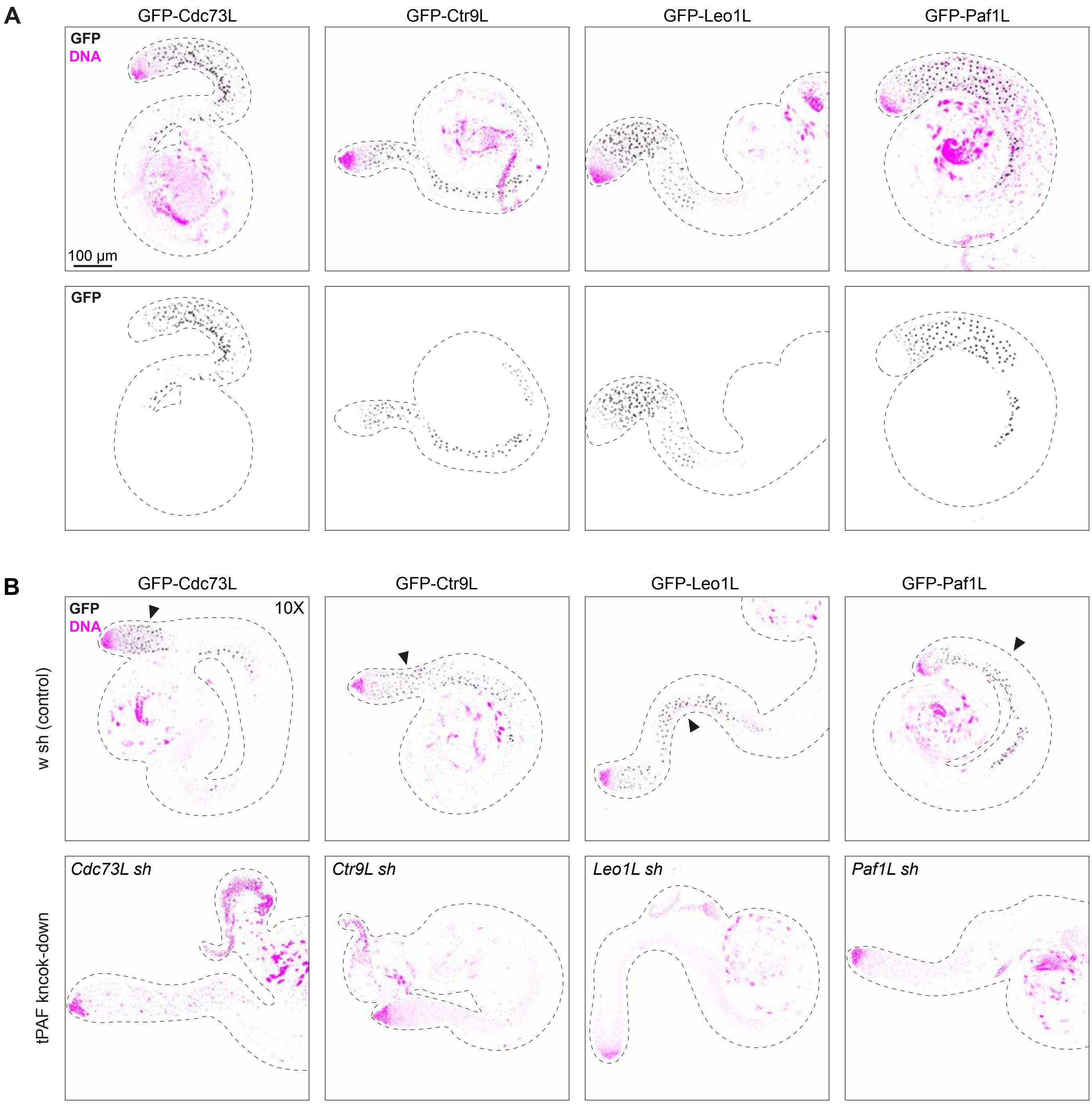


**FIGURE S7**

(A) Confocal images showing the GFP fluorescence signal in testis tissue from the endogenously tagged (3xFLAG-3xV5-GFP-)tPAF proteins noted above the images. Upper images show the GFP signal merged with DAPI-stained DNA shown in magenta. The lower images show the GFP channel signal only. (B) As (A) but with upper images deriving from control germline knockdown flies (w sh) and the lower images from flies subjected to knockdown targeting the tagged tPAF gene as noted in the upper left corner of each image.

#### Figure S8

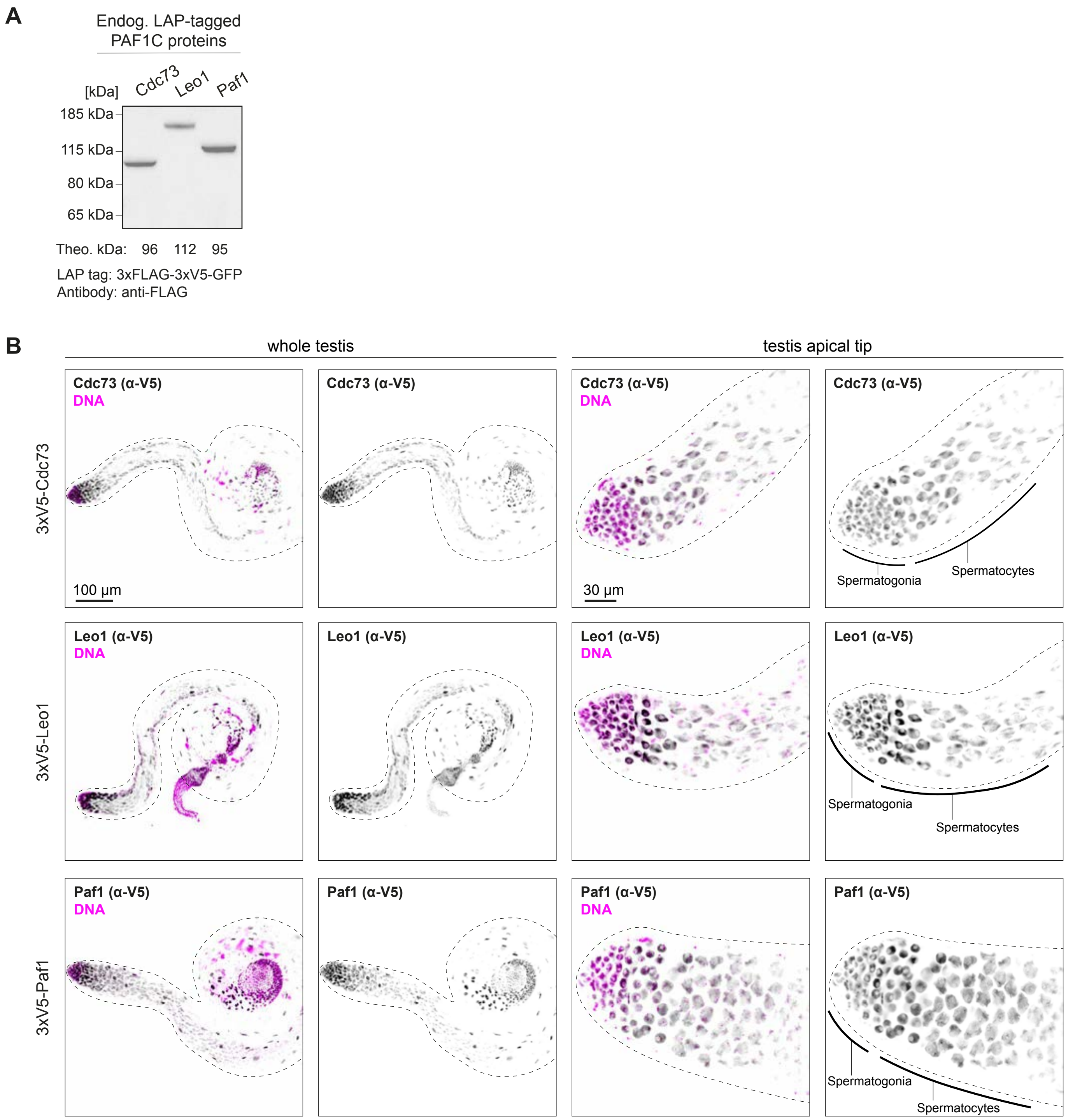


**FIGURE S8**
(A) Western blot detection of endogenously tagged (3xFLAG-3xV5-GFP-)PAF1C proteins using anti-FLAG antibodies. (B) Confocal images showing the anti-V5 immunofluorescence signal in either whole testis tissue (left half) or in the testis apical tip (right half) from the endogenously tagged (3xFLAG-3xV5-GFP-)PAF1C proteins noted in the images. The leftmost images show the GFP signal merged with DAPI-stained DNA shown in magenta. The rightmost images show the GFP channel signal only.

**Figure S9**

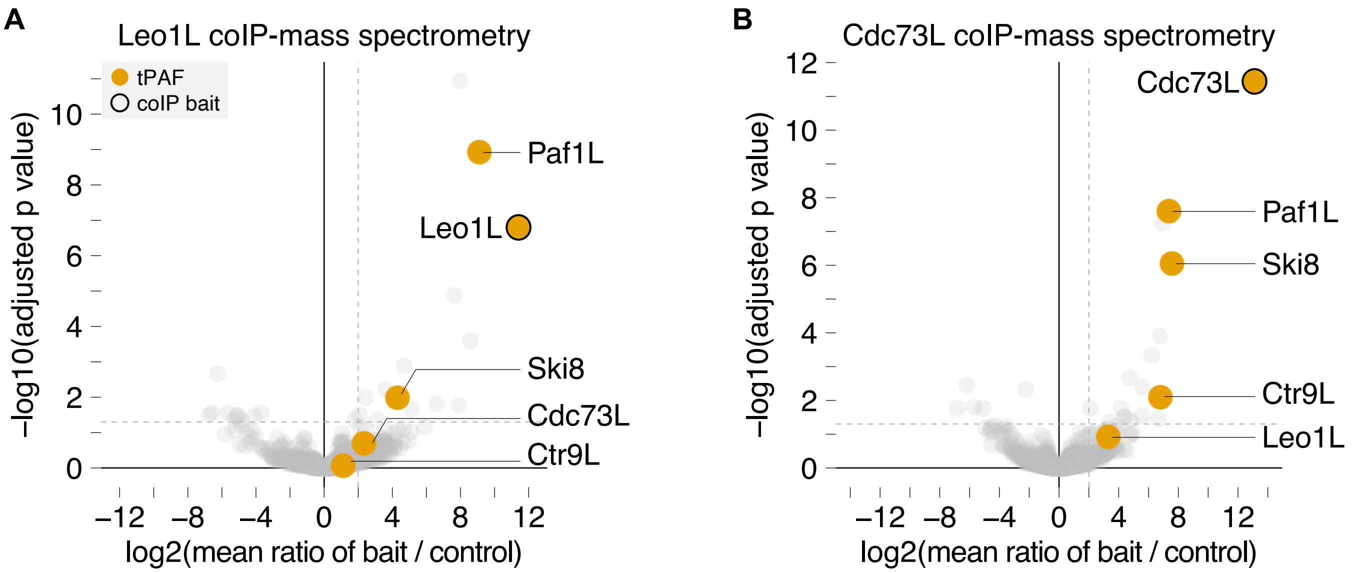


**FIGURE S9**
(A-B) Enrichment values and corresponding statistical significance levels for proteins co-immunopurified (coIP) with endogenously tagged Leo1L (A) or Cdc73L (B) proteins as detected by mass-spectrometry (biological replicates, n, are indicated in Figure 3F)

#### Figure S10

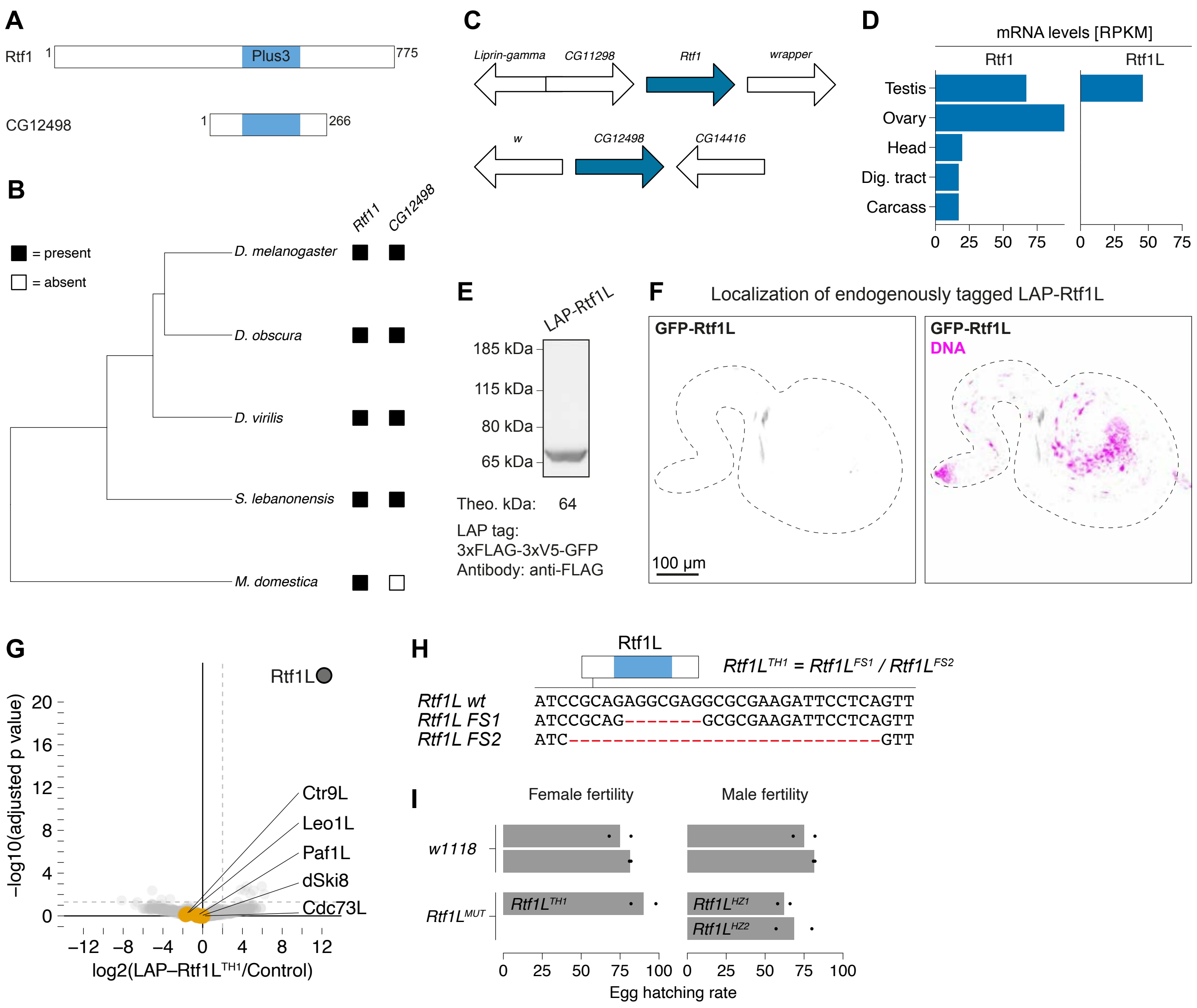


**FIGURE S10. Rtf1L (CG12498) is expressed in Drosophila melanogaster testis but shows no evidence of physical or functional interaction with tPAF**

(A) Schematic of the aligned canonical Rtf1 and Rtf1L (CG12498) genes with the main protein domains indicated. (B) Phylogenetic tree of the indicated species (left) with schematic indication of the presence (solid black box) or absence (white box) of the Rtf1 and Rtf1L genes as assessed by BLAST search and synteny analysis. (C) Schematic representation of the gene annotation context of the Rtf1 and Rtf1L genes used for synteny analysis. (D) mRNA levels of the Rtf1 and Rtf1L genes in selected adult tissues (RNAseq data from modENCODE, ref. 83), RPKM = Reads Per Kilobase per Million mapped reads. (E) Western blot detection of endogenously tagged (3xFLAG-3xV5-GFP-)Rtf1L proteins using anti-FLAG antibodies. (F) Confocal images showing the GFP signal in testis tissue from endogenously tagged (3xFLAG-3xV5-GFP-)Rtf1L proteins with DAPI-stained DNA are shown in magenta. (G) Enrichment values and corresponding statistical significance levels for proteins co-immunopurified (coIP) with endogenously tagged Rtf1L proteins as detected by mass-spectrometry. (H) Schematic showing Rtf1L with protein domains from (A) indicated in blue. The nucleotide sequence changes in the CRISPR/Cas9-generated Rtf1L mutant alleles are illustrated below the gene schematics. (I) Bar plot showing the mean embryo hatching rate in percent of eggs laid by control (*w*$^{1118}$) and Rtf1L mutant flies. Individual data points from biological replicates (n = 2) are shown as black dots. HZ1 = hemizygous Rtf1LFS1 mutants. HZ2 = hemizygous *Rtf1L*$^{FS2}$ mutants. TH1 = transheterozygous *Rtf1L*$^{FS1}$ / *Rtf1L*$^{FS2}$ mutants.

#### Figure S11

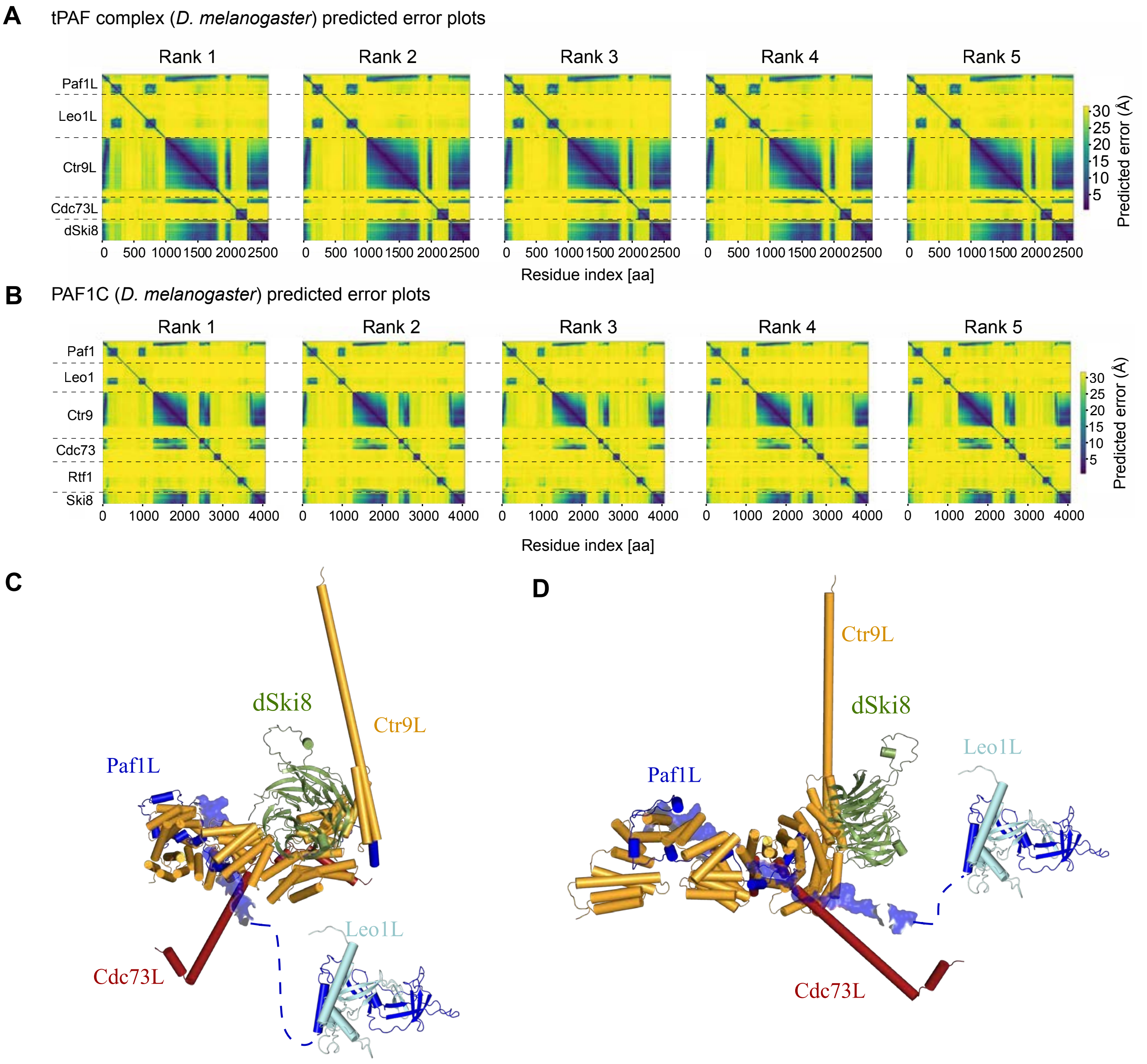


**FIGURE S11.**
(A-B) Heat maps showing predicted error (Å) values of five Alphafold2 models made for the *Drosophila melanogaster* tPAF (A) and PAF1C (B) complexes. Protein identities are indicated on the y axis and the corresponding protein complex concatenated amino acid count is listed on the x axis. (C-D) Additional Alphafold2-predicted structure models of the tPAF complex that differ from the model shown in Figure 3G, mainly by the positioning of the Paf1L protein, which influences the predicted position of the Paf1L/Leo1L subdomain. Dashed lines indicate manually positioned structured domains.

#### Figure S12

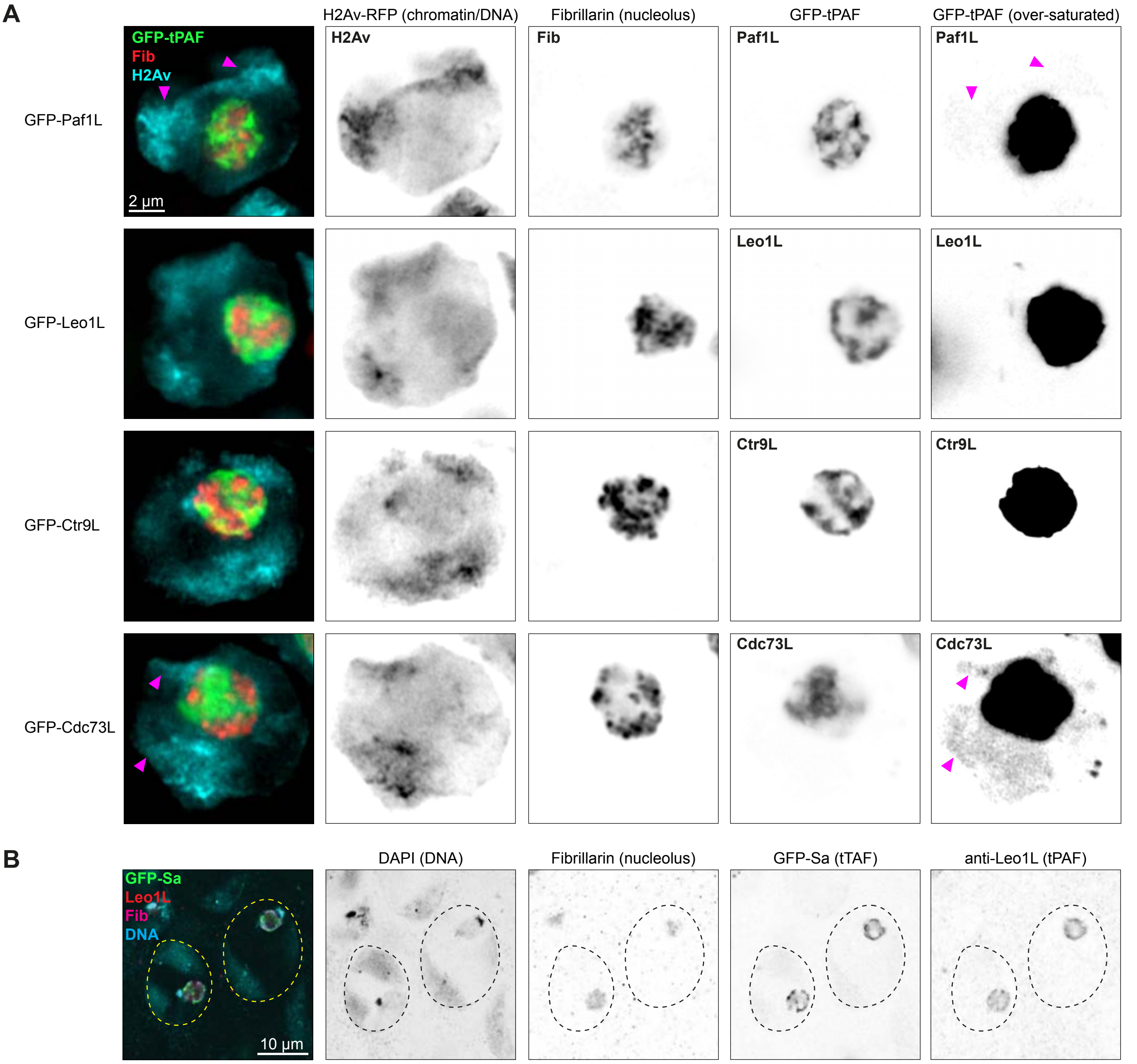


**FIGURE S12.**
(A) Confocal microscopy images of testis spermatocytes nuclei showing the localization of endogenously tagged (3xFLAG-3xV5-GFP)-tPAF proteins, Fibrillarin (Fib, anti-Fib immunoflourence staining) and chromatin visualized by H2Av-RFP tagged histone proteins. The dashed magenta line indicates the border of the nucleo. Magenta arrowheads indicate the position of major chromosome territories. (B) Confocal microscopy images showing the localization of GFP-Sa (tTAF) proteins, Leo1L (tPAF, anti-Leo1L immunoflourence staining), (Fibrillarin (Fib, anti-Fib immunoflourence staining) and chromatin visualized by DNA staining using DAPI.

### Figure S13

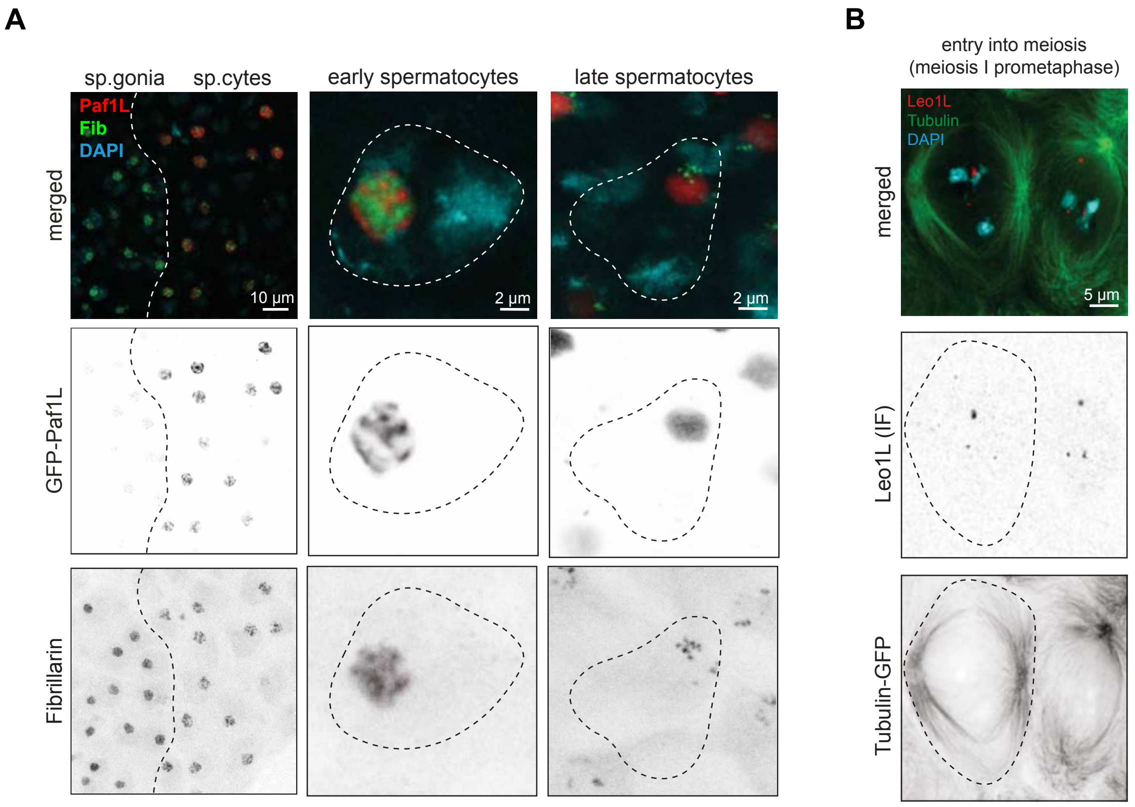


**FIGURE S13.**
(A) Confocal microscopy images showing the localization of endogenously tagged (3xFLAG-3xV5-GFP)-Paf1L (tPAF) proteins, (Fibrillarin (Fib, anti-Fib immunoflourence staining) and chromatin visualized by DNA staining using DAPI. The images show localization patterns through the progression of spermatocyte development with spermatogonia to spermatocyte transition (leftmost images), early spermatocytes (middle), and late spermatocytes (rightmost images). (B) Confocal microscopy images of spermatocyte cells in meiosis I prometaphase showing the localization of Leo1L (tPAF, anti-Leo1L immunoflourence staining), Tubulin (anti-Tubulin immunoflourence staining) and DNA staining using DAPI.

#### Figure S14

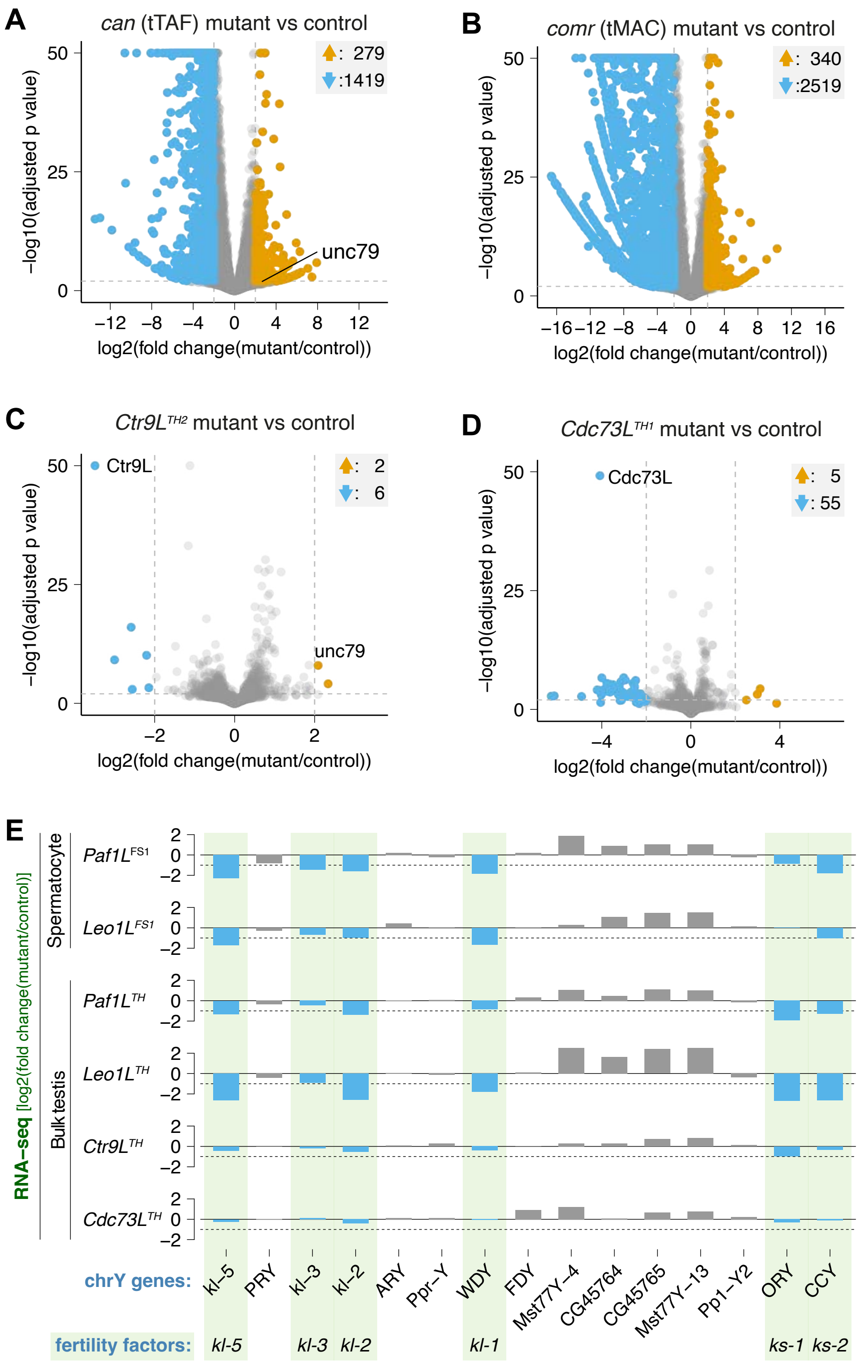


**FIGURE S14.**
(A-D) Enrichment values and corresponding statistical significance levels for mRNAs quantification based on bulk testis RNAseq in biological replicates of (A) *can* (tTAF), (B) *comr* (tMAC), (C) *Ctr9L* (tPAF), and (D) *Cdc73L* (tPAF) mutants compared to control samples. Orange and blue arrows indicate the number of genes categorized as upregulated and downregulated, respectively. The can and comr mutant data (A-B) is derived from reference 49. (E) Bar plots showing the log2 fold change in mRNA levels of Y-linked genes as measured by RNA-seq, comparing tPAF mutant and control genotypes. Blue bars: mRNA level at least 50 % decreased in any tPAF mutant sample compared to control (Figure S14E). Green highlights: male fertility factor genes.

#### Figure S15

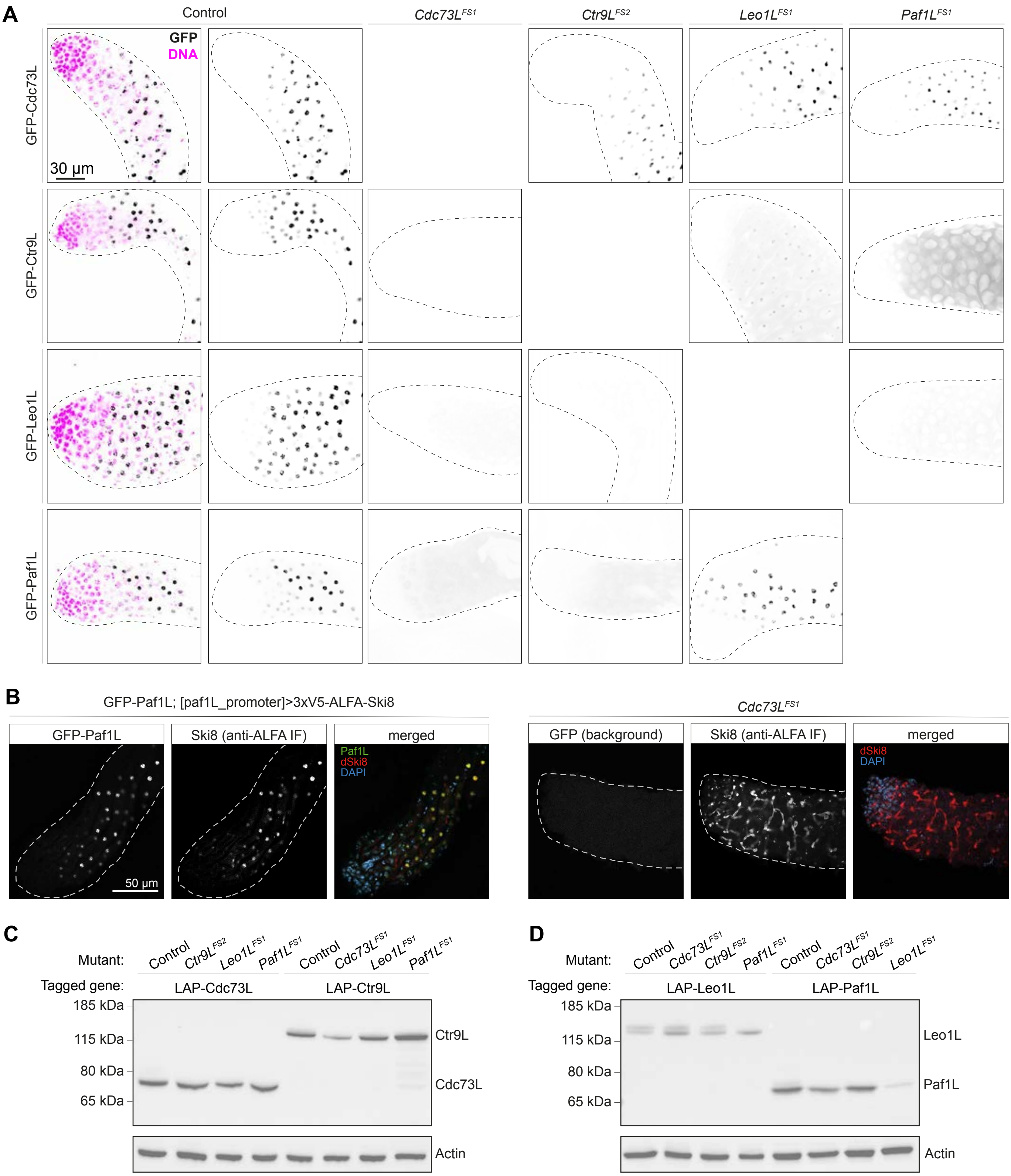


**Figure S15.** See next page for legend

#### Figure S16

**FIGURE S15.** (continued from last page)
(A) Confocal images showing the GFP signal in testis apical tip tissue from endogenously tagged (3xFLAG-3xV5-GFP)-tPAF proteins with DAPI-stained DNA shown in magenta. The localization of each core tPAF protein (see annotations to the left) are shown in control (two leftmost columns) or mutant background for the other tPAF core genes. The dashed line indicates the tissue outline based on DNA DAPI staining. (B) Confocal microscopy images showing the localization of endogenously tagged (3xFLAG-3xV5-GFP)-Paf1L (tPAF) proteins, together with tagged (3xV5-ALFA)-Ski8 protein expressed under the control of the Paf1L promoter as well as DAPI-stained DNA shown in blue. (C-D) Western blot detection of endogenously tagged tPAF proteins in control or mutant background for the other tPAF core genes. Tagged proteins were detected using anti-FLAG antibodies and anti-Actin antibodies were used for the Actin loading control.

---

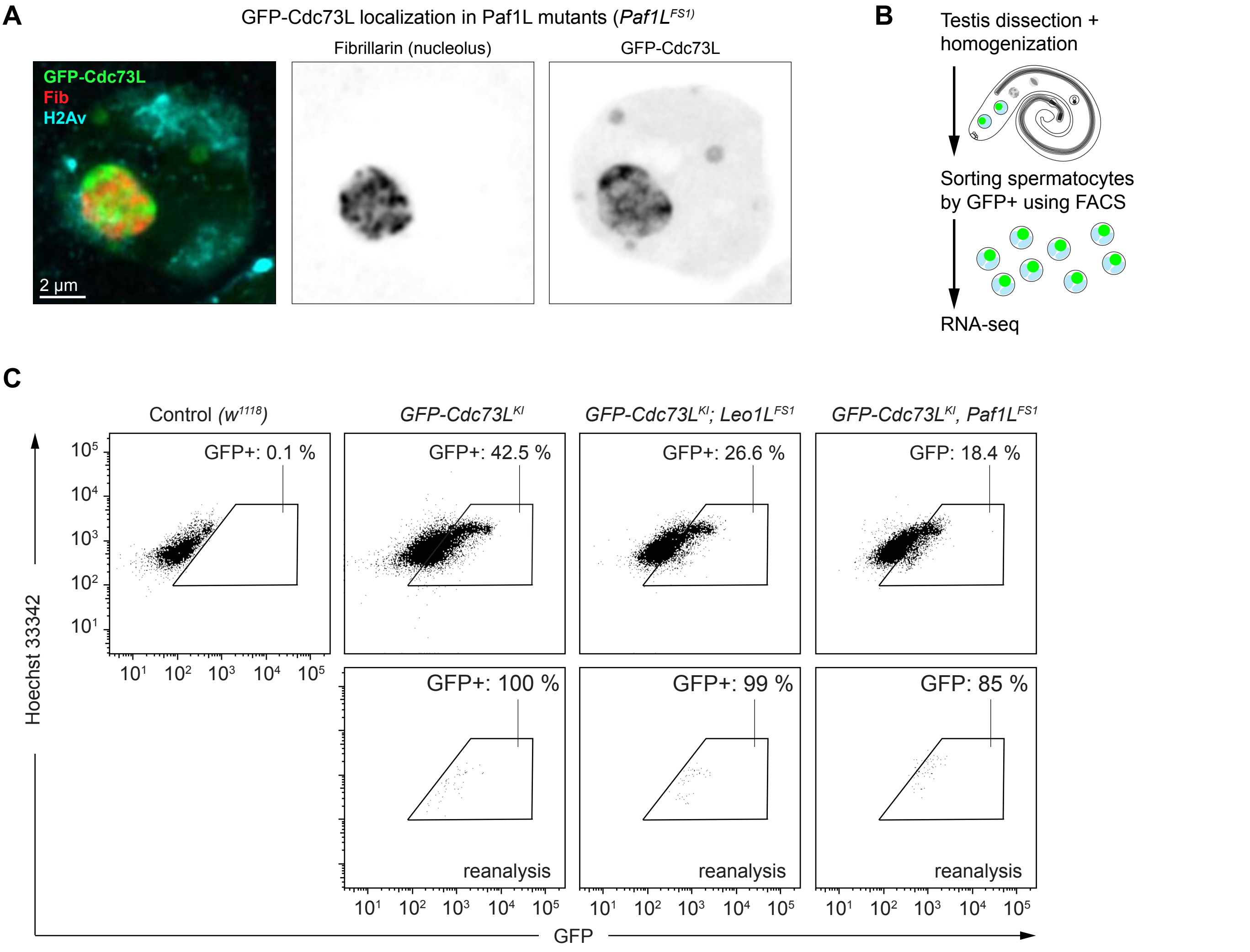


**FIGURE S16.**
(A) Confocal microscopy images of testis spermatocytes nuclei in *Paf1L* mutant background. The images show the localization of endogenously tagged (3xFLAG-3xV5-GFP)-Cdc73L (tPAF) proteins, Fibrillarin (Fib, anti-Fib immunoflourence staining) and chromatin visualized by H2Av-RFP tagged histone proteins. (B) Schematic illustrating the experimental strategy for spermatocyte transcriptome analyses. (C) FACS scatter plots showing GFP (x-axis) and DNA Hoechst (y-axis) intensities of the detected cells with the GFP+ gate used for spermatocyte sorting outline with a black line. The top plots show data collected during primary sorting and the bottom plots show the re-analysis to verify the sorted samples. Control genotype testis served as a control to set the gate for GFP-expressing cells.

Figure S17

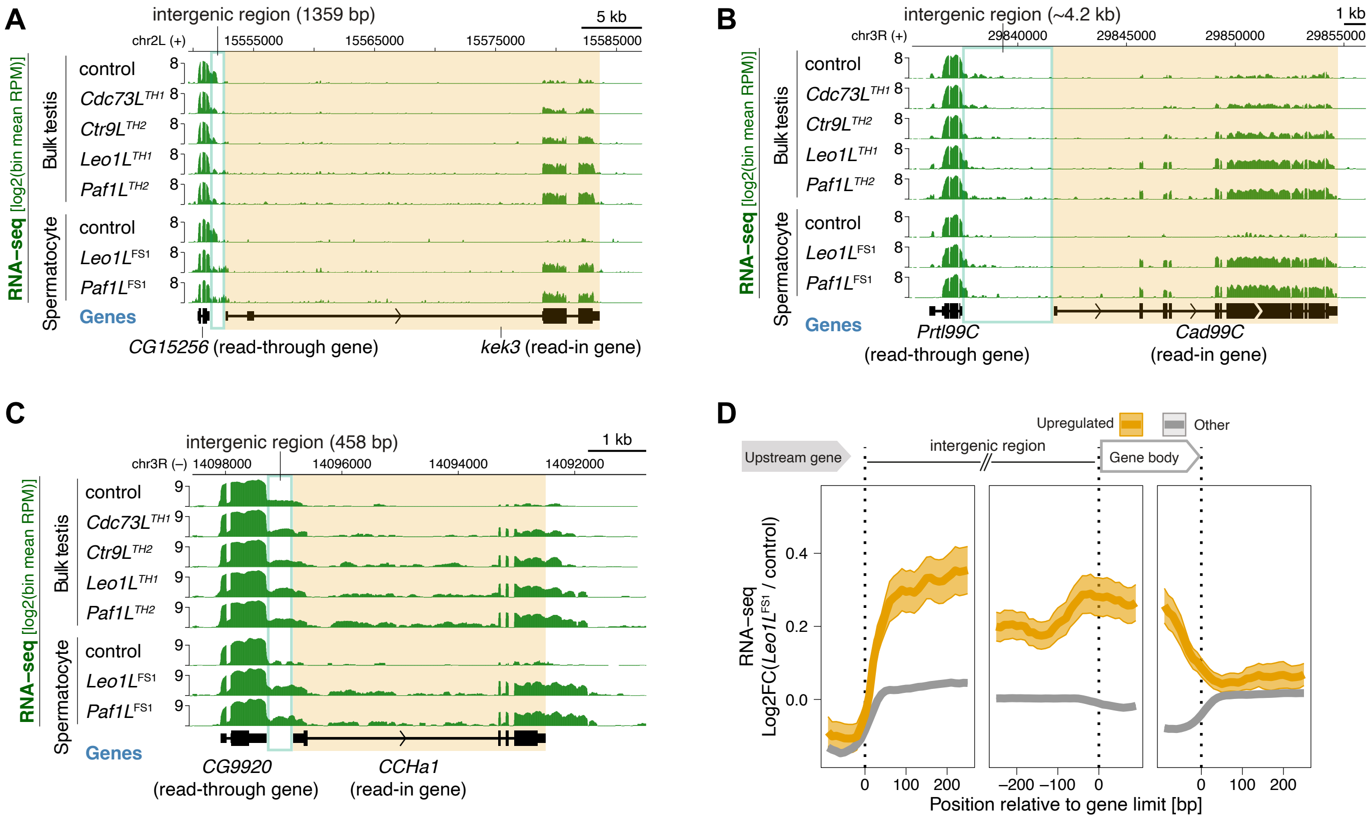


**FIGURE S17.**
(A-C) Genome tracks showing RNA-seq coverage around various genes upregulated in tPAF mutants for the indicated genotypes and sample type. The coverage signal is displayed as the log2 of mean reads per million mapped reads (RPM) per genome region bin. (D) Metagene analysis of RNAseq coverage fold change in *Leo1L* mutants relative to control samples. Standard error is plotted as a faded ribbon around the solid line showing the mean value at each genomic position. The plots compare coverage around bulk annotated genes ("other") to that of genes upregulated in *Leo1L* tPAF mutants.

#### Figure S18

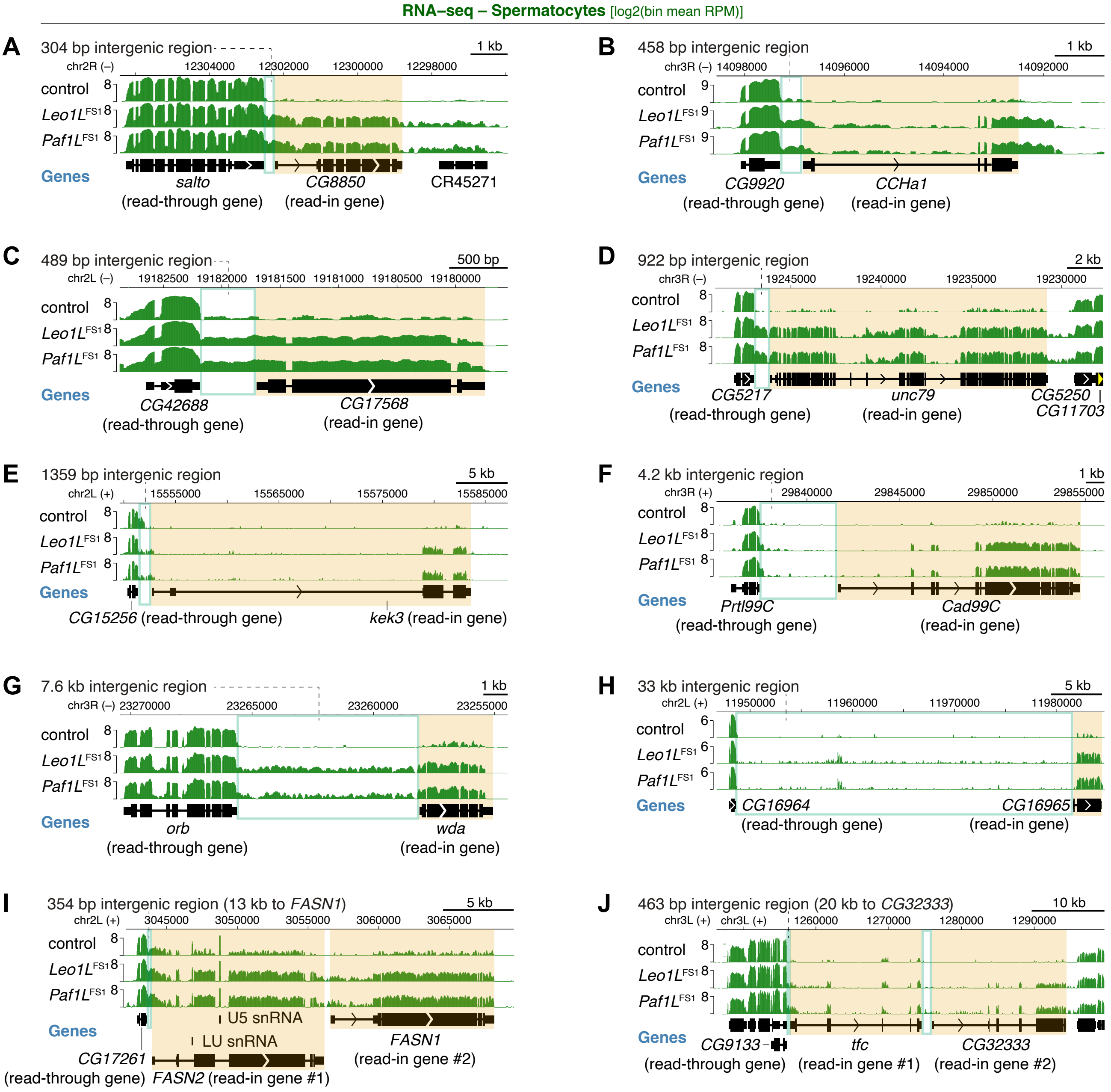


**FIGURE S18.**
(A-J) Genome tracks showing RNA-seq coverage around various genes upregulated in tPAF mutants for the indicated genotypes and sample type. The coverage signal is displayed as the log2 of mean reads per million mapped reads (RPM) per genome region bin.

#### Figure S19

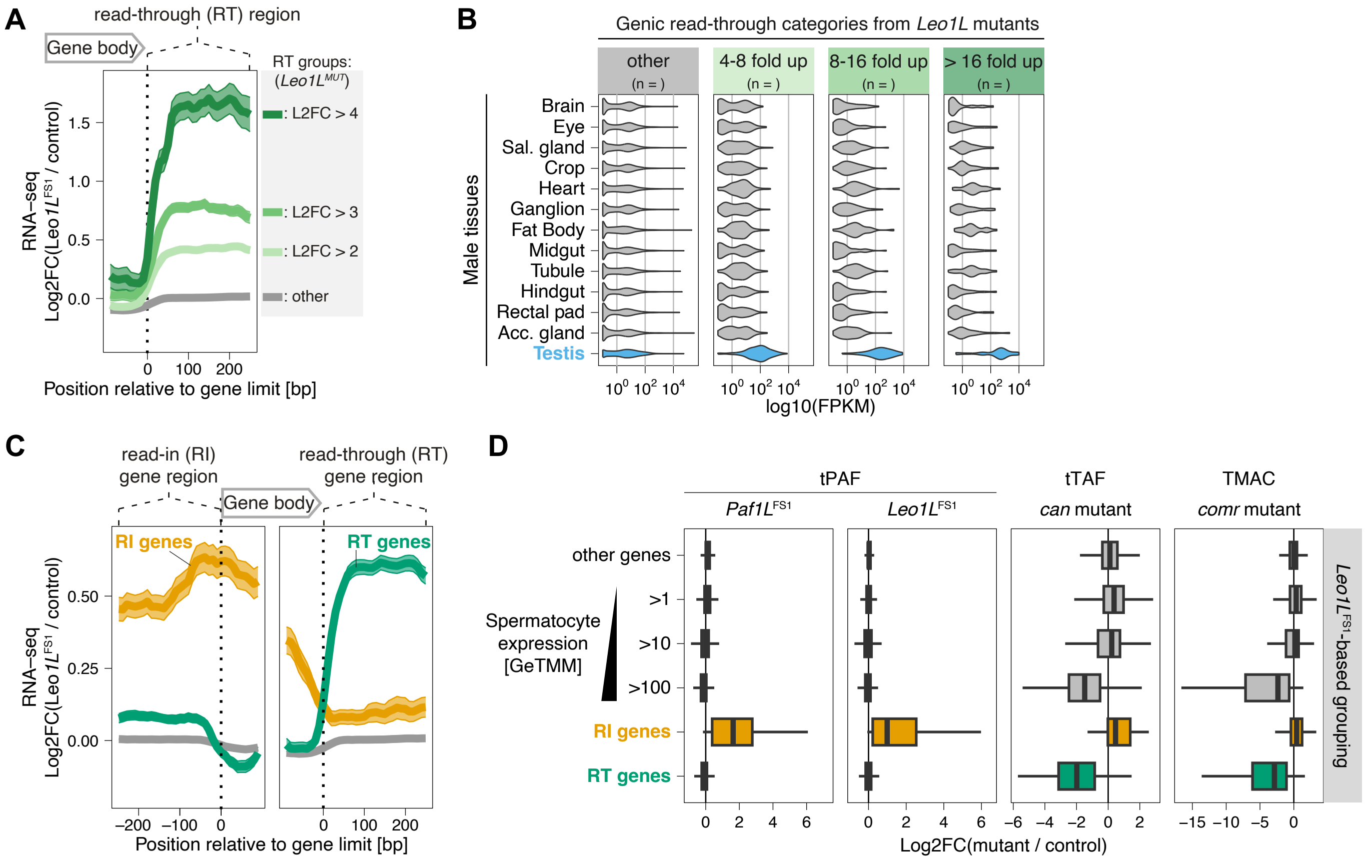


**FIGURE S19.**
(A) Metagene analysis of RNAseq coverage fold change in *Leo1L* mutants relative to control samples. Standard error is plotted as a faded ribbon around the a solid line showing the mean value at each genomic position. The plots compare coverage around bulk annotated genes ("other") to that of genes showing increased downstream RNA-seq signal in *Leo1L* tPAF mutants. (B) Violin plot of gene expression levels based on FlyAtlas2 RNAseq from the listed fly tissues. Expression is plotted separately for the gene categories established in (A). FPKM = Fragments Per Kilobase per Million mapped fragments. (C) Like (A) but here comparing read-in (RI) and read-through (RT) gene categories to the remaining ("other") annotated gene loci. (D) Box plots showing the distribution of change in gene expression values between mutant and control RNA-seq samples for the listed gene categories (based on C) and mutant genotypes. The data in the tPAF mutant panels is the spermatocyte RNA-seq data. The data underlying the tTAF and tMAC panels originates from reference 44.

Figure S20

A

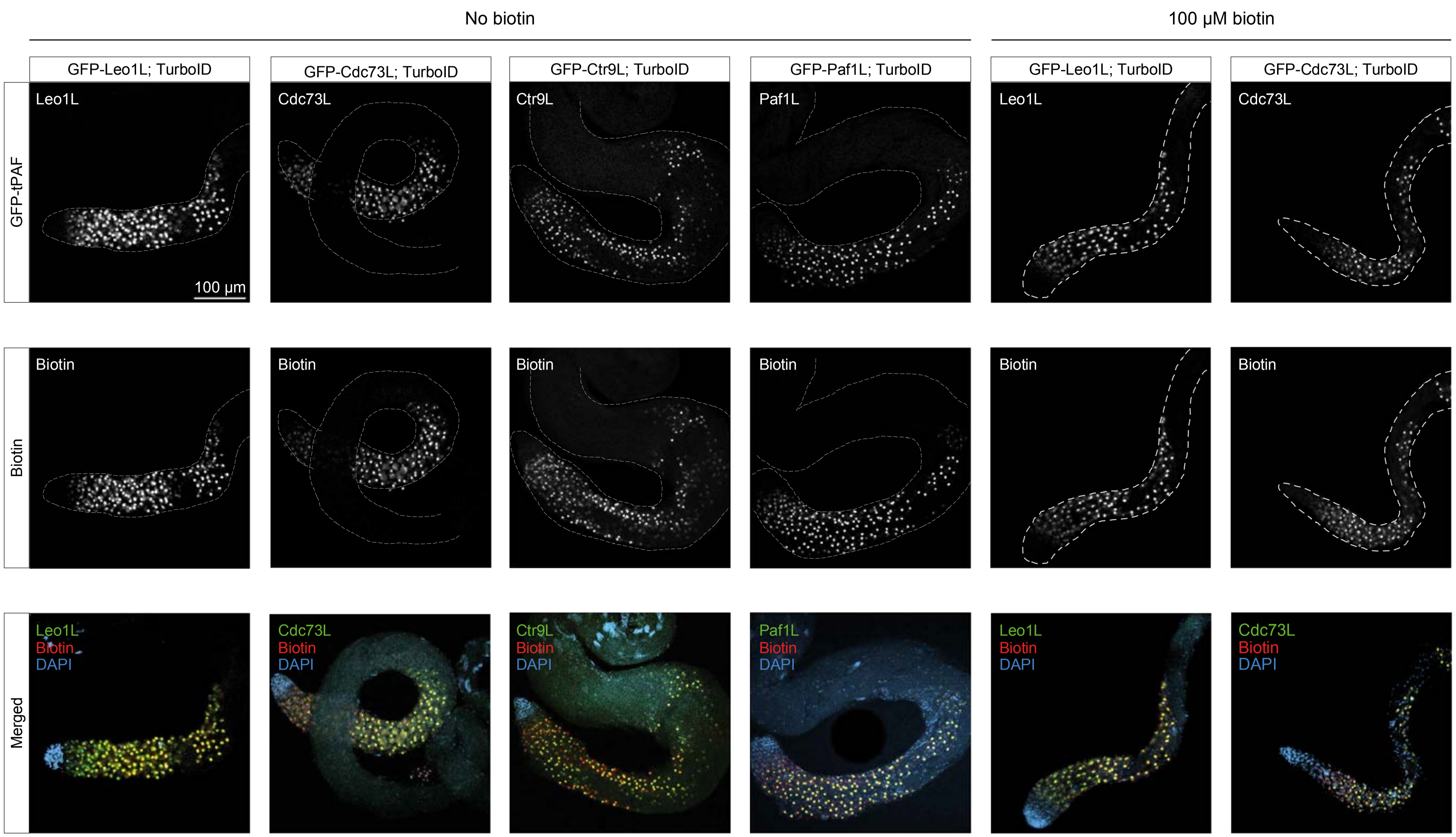


**Figure S20.**
(A) Confocal microscopy images showing whole testis tissue from flies harboring endogenously tagged (3xFLAG-3xV5-GFP)-tPAF proteins combined with a TurboID cassette. The images display localization of GFP signal from tagged tPAF proteins, biotin accumulation (Streptavidin-AlexoFluor647), and chromatin visualized by DNA DAPI staining. The dashed lines indicate the border of the testis tissue. The rightmost images show tissue from flies reared on food containing 100 µM biotin supplement.

#### Figure S21

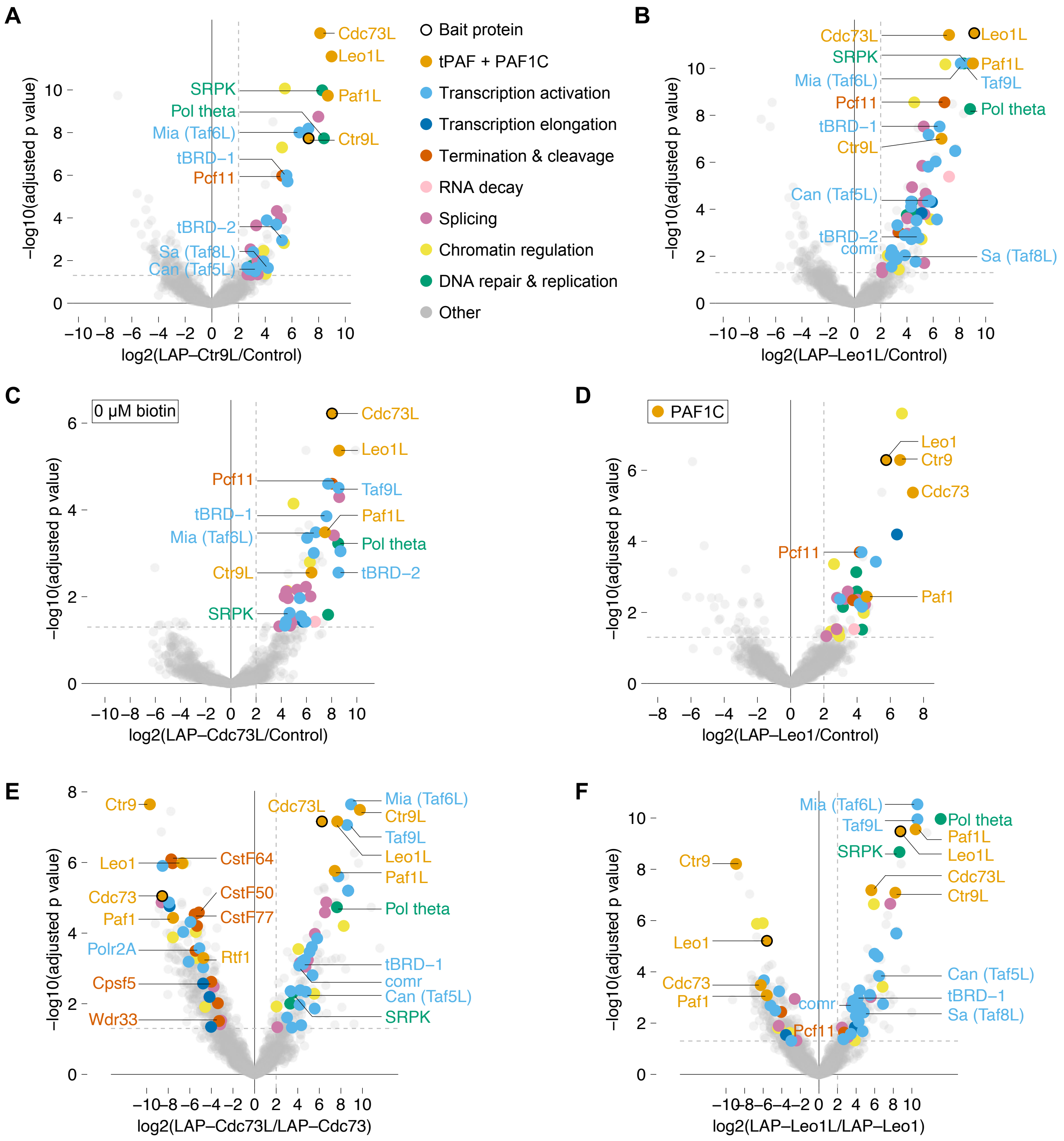


**FIGURE S21.**
(A-F) Enrichment values and corresponding statistical significance levels for proteins associated with the proximity labeling bait protein compared to control samples (except E-F) as detected by proximity labeling mass-spectrometry. The baits for the displayed data are Ctr9L (A), Leo1L (B), Cdc73L without biotin supplement (C), and Leo1 (PAF1C) (D). (E-F) directly compare detected protein abundances between tPAF and PAF1C paralog baits with Cdc73L and Cdc73 being compared in (E) and Leo1L and Leo1 in (F). The colored highlights indicate manually annotated protein groups as listed to the right of (A).

#### Figure S22

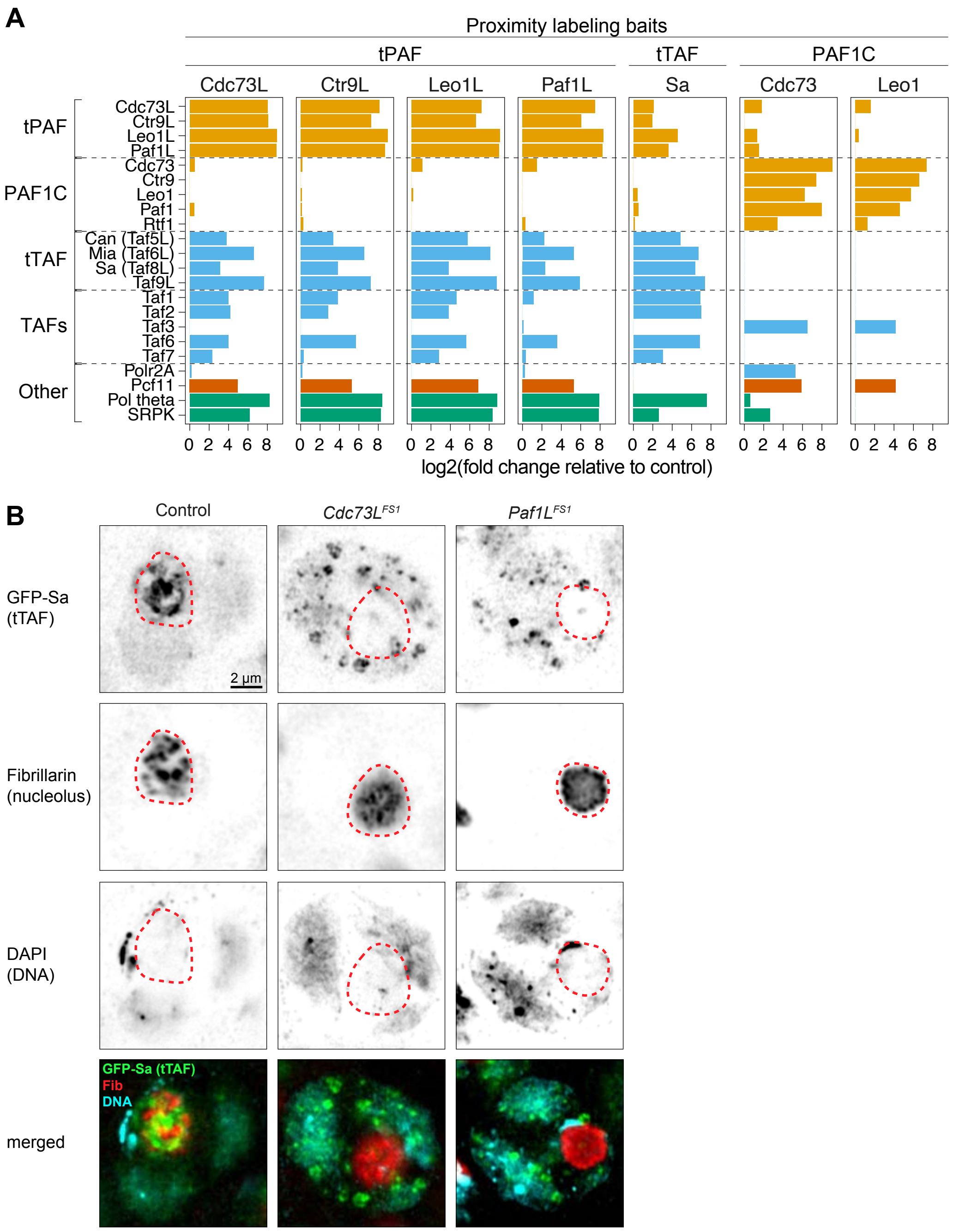


**FIGURE S22.**
(A) Bar plot showing a comparison of enrichment for select proteins in different bait protein samples. Proximity labing samples from tPAF, tTAF, and PAF1C bait proteins are compared as shown. (B) Confocal microscopy images showing the localization of GFP-tagged Sa (tTAF), Fibrillarin (Fib, anti-Fib immunoflourence staining), and chromatin visualized by DAPI. Control ($w^{1118}$) genotype as well as *Cdc73L* and *Paf1L* mutant are shown. Dashed red line: nucleolus border.
